# Allelic Variation of *MYB10* is the Major Force Controlling Natural Variation of Skin and Flesh Color in Strawberry (*Fragaria* spp.) fruit

**DOI:** 10.1101/2020.06.12.148015

**Authors:** Cristina Castillejo, Veronika Waurich, Henning Wagner, Rubén Ramos, Nicolás Oiza, Pilar Muñoz, Juan C. Triviño, Julie Caruana, Zhongchi Liu, Nicolás Cobo, Michael A. Hardigan, Steven J. Knapp, José G. Vallarino, Sonia Osorio, Carmen Martín-Pizarro, David Posé, Tuomas Toivainen, Timo Hytönen, Youngjae Oh, Christopher R. Barbey, Vance M. Whitaker, Seonghee Lee, Klaus Olbricht, José F. Sánchez-Sevilla, Iraida Amaya

**Author notes:** Corresponding author: Iraida Amaya.

## Abstract

Anthocyanins are the principal color-producing compounds synthesized in developing fruits of strawberry (*Fragaria* spp.). Substantial natural variation in color have been observed in fruits of diploid and octoploid accessions, resulting from distinct accumulation and distribution of anthocyanins in fruits. Anthocyanin biosynthesis is controlled by a clade of R2R3 MYB transcription factors, among which *MYB10* has been shown as the main activator in strawberry fruit. Here, we show that *MYB10* mutations cause most of the anthocyanin variation observed in diploid woodland strawberry *(F. vesca)* and octoploid cultivated strawberry (*F*. ×*ananassa*). Using a mapping-by-sequencing approach, we identified a *gypsy*-transposon insertion in *MYB10* that truncates the protein and knocks out anthocyanin biosynthesis in a white-fruited *F. vesca* ecotype. Two additional loss-of-function *MYB10* mutations were identified among geographically diverse white-fruited *F. vesca* ecotypes. Genetic and transcriptomic analyses in octoploid *Fragaria spp.* revealed that *FaMYB10-2,* one of three *MYB10* homoeologs identified, residing in the *F. iinumae-derived* subgenome, regulates the biosynthesis of anthocyanins in developing fruit. Furthermore, independent mutations in *MYB10-2* are the underlying cause of natural variation in fruit skin and flesh color in octoploid strawberry. We identified a CACTA-like transposon *(FaEnSpm-2)* insertion in the *MYB10-2* promoter of red-fleshed accessions that was associated with enhanced expression and anthocyanin accumulation. Our findings suggest that putative cis regulatory elements provided by *FaEnSpm-2* are required for high and ectopic *MYB10-2* expression and induction of anthocyanin biosynthesis in fruit flesh. We developed *MYB10-2* (sub-genome) specific DNA markers for marker-assisted selection that accurately predicted anthocyanin phenotypes in octoploid segregating populations.

## INTRODUCTION

The characteristic red color of strawberry fruit is due to the accumulation of anthocyanins, water-soluble pigments synthesized through the flavonoid pathway (Almeida et al., 2007; Tohge et al., 2017). The initial substrate, 4-coumaroyl-coenzyme A (CoA,) is produced from phenylalanine through the sequential activity of the general phenylpropanoid pathway enzymes phenylalanine ammonia-lyase (PAL), cinnamate 4-hydroxylase (C4H) and 4-coumarate:coenzyme A ligase (4CL). The flavonoid pathway begins with the condensation of one molecule of 4-coumaroyl-CoA and three molecules of malonyl-CoA by chalcone synthase (CHS). Chalcone isomerase (CHI) subsequently converts naringenin chalcone into naringenin. Following steps involving flavonoid 3-hydroxylase (F3H) and flavonoid 3’,5’-hydroxylase (F3’5’H) generate dihydroflavonols. Genes encoding enzymes involved in these steps have been named early biosynthetic genes (EBGs). Downstream genes of the pathway are usually named late biosynthetic genes (LBGs) and encode for dihydroflavonol 4-reductase (DFR) that generate colorless leucoanthocyanidins, anthocyanidin synthase (ANS) producing anthocyanidins, and several glycosyltransferases (UFGTs) that attach sugar molecules to anthocyanidins to generate the first stable anthocyanins. Anthocyanins are synthesized at the endoplasmic reticulum and are later transported into the vacuole for storage via different types of mechanisms including glutathione S-transferases (GSTs; Luo et al., 2018). Anthocyanin biosynthesis is tuned through transcriptional regulation of structural genes by transcription factors that include MYB, bHLH, and WD-repeat proteins associated in what is called the MBW ternary complex (Xu et al., 2015; Zhang et al., 2014; Jaakola, 2013). MYB transcription factors are often the major determinant of natural variation in anthocyanin biosynthesis (Allan et al., 2008).

Fruit color is a key quality trait for fruit breeders. Fruit color in the genus *Fragaria* varies widely from completely white fruits to dark red, with wide variation in the internal concentration and distribution of anthocyanins throughout the fruit (Hancock, 1999; Hancock et al., 2003). Gaining insight into the genetic factors affecting natural variation in external (skin) and internal (flesh) fruit color is key for efficient modification of this trait in new cultivars and will facilitate the rapid development of fruits with increased or reduced levels of anthocyanins. Besides contributing to fruit color, anthocyanins possess anti-oxidative properties that have been associated with several health-promoting effects and positive impacts on cardiovascular disorders and degenerative diseases (He and Giusti, 2010; Forbes-Hernandez et al., 2016; Del Rio et al., 2013).

The genus *Fragaria* belongs to the Rosaceae family and comprises 24 species, including the world-wide cultivated strawberry *(Fragaria ×ananassa;* Liston et al., 2014; Staudt, 2009). The genus displays a series of ploidy levels, ranging from diploid species such as *Fragaria vesca* (2n=2x=14) to decaploid species such as some accessions of *Fragaria iturupensis* (2n=10x=70). The cultivated strawberry, *Fragaria* ×*ananassa*, originated in France nearly 300 years ago via hybridization between two wild octoploid species, *Fragaria chiloensis* and *Fragaria virginiana,* which were introduced from South and North America, respectively (Darrow, 1966; Hancock, 1999). Cultivated strawberry and its wild progenitor species are allopolyploids with 2n=8x=56 chromosomes. Recent studies agree that one of the four subgenomes of the octoploid *Fragaria* species originates from a *F. vesca* ancestor and another from a *F. iinumae* ancestor, while the origin of the remaining two subgenomes, possibly related to the extant species *Fragaria viridis* and *Fragaria nipponica,* is still under investigation (Tennessen et al., 2014; Edger et al., 2019a; Liston et al., 2019; Edger et al., 2019b).

Due to its simpler diploid genome, *F. vesca* is a model for genetic and genomic studies, and efficient genomic resources have been generated including transcriptomes (Hollender et al., 2012; 2014; Li et al., 2019) and a recently improved near-complete genome sequence (Shulaev et al., 2011; Edger et al., 2018). *F. vesca* accessions with white fruits, including the sequenced Hawaii4 accession, have been described and stored in multiple germplasm repositories. Despite having red skin color, *F. vesca* accessions, such as ‘Reine des Vallées’ or ‘Ruegen’, are all characterized by white or pale-yellow flesh (NCGR, Corvallis repository). Red versus white external fruit color in *F. vesca* is governed by a single locus named *C* (Brown and Wareing, 1965). The *c* locus from the ‘Yellow Wonder’ cultivar was subsequently mapped at the bottom of linkage group 1 (Williamson et al., 1995; Deng and Davis, 2001). Recently, a genome-scale variant analysis showed that a single nucleotide polymorphism (SNP) (G36C) causing an amino acid change (W12S) in the *FvMYB10* gene was responsible for the loss of anthocyanins and pale color of ‘Yellow Wonder’ fruits (Hawkins et al., 2016). Several studies have addressed the characterization of most of the structural and regulatory genes necessary for color development in the genus *Fragaria* (Lunkenbein et al., 2006; Carbone et al., 2009; Hossain et al., 2018; Miosic et al., 2014; Lin-Wang et al., 2014; Salvatierra et al., 2010; 2013; Medina-Puche et al., 2014; Fischer et al., 2014; Duan et al., 2017; Thill et al., 2013; Griesser et al., 2008; Moyano et al., 1998; and references therein). The identification of genes or genomic regions governing fruit color variation in cultivated strawberry is crucial for hastening the development of new cultivars with desired characteristics. However, the octoploid nature of this species complicates genetic analyses because up to eight alleles can be found if each homoeologous locus from the four sub-genomes were conserved after polyploidization (Edger et al., 2019a). A number of studies have detected quantitative trait loci (QTL) contributing to external color intensity or anthocyanin content of strawberry fruits (Zorrilla-Fontanesi et al., 2011; Lerceteau-Kohler et al., 2012; Castro and Lewers, 2016). Each of these three studies have identified three to six genomic regions contributing to color-related traits, but the phenotypic variation explained by each QTL was relatively low, with only few QTLs explaining more than 15% of phenotypic variation. More recently, an 8bp insertion in the coding region of *FaMYB10* has been associated with loss of anthocyanins in fruits of ‘Snow Princess’, an octoploid strawberry cultivar with completely white fruits (Wang et al., 2020), but its subgenome location was not described. Three full-length homoeologous *FaMYB10* genes were annotated in the octoploid ‘Camarosa’ genome (Edger et al., 2019a). The patterns of expression and roles of these genes in determining fruit color variation are not fully known. Are all three genes mutated in white octoploid strawberry? How many homoeologs are expressed in developing fruit?

In this work, we have carried out an extensive phenotypic, genetic and molecular analysis of *Fragaria* genetic resources to identify genetic determinants contributing to natural fruit color variation in strawberry. First, we screened available diversity for fruit color in the diploid *F. vesca* and identified novel mutations for the loss of anthocyanins and associated red color. We then extended the analysis to seven octoploid accessions with fruit with either white flesh/red skin or white flesh/white skin, in comparison to fruits with red flesh/red skin. One of our objectives was to identify specific alleles in cultivated strawberry that could be targeted for marker development that would aid breeders in the efficient development of new improved cultivars with desired fruit color. Strikingly, our results show that all analyzed color variants in the genus *Fragaria* are caused by independent mutations in the same gene, the transcription factor *MYB10.* Different genetic lesions are responsible for distinct flesh and skin color phenotypes, and therefore, allele-specific markers will need to be developed to track specific traits. We further show that independent mutations in only one of three *MYB10* homoeologs *(MYB10-2)* cause skin- and flesh-color variation in octoploid strawberry. Red-flesh color was linked to a CACTA-like transposon insertion in the *FaMYB10* promoter while white-flesh mutant alleles were inherited from white-fleshed *F. chiloensis* donors. We first developed a high-resolution melting (HRM) marker able to predict white fruit skin color derived from a mutation in the coding region of *FaMYB10.* Subsequently, two additional DNA markers (one based on PCR and agarose gel and another on Kompetitive allele-specific PCR) were developed that predict strawberry internal fruit color in diverse octoploid germplasm. The described markers represent useful tools for selection of fruit color, particularly when *F. chiloensis* accessions are used in breeding programs.

## RESULTS AND DISCUSSION

### Identification of loci controlling external fruit color in *F. vesca*

To identify genetic factors affecting fruit color in *F. vesca,* we developed an F_2_ mapping population of 145 lines derived from ‘Reine des Vallées’ (RV660) and the IFAPA white-fruited accession ESP138.596 (WV596). As previously described for other backgrounds (Brown and Wareing, 1965), red vs. white fruit color in the RV660 × WV596 F_2_ population segregated as expected for a single mutation (1:3; χ^2^ test, *p* = 0.922). In order to fast-map the locus controlling red color in this *F. vesca* population, two pools were prepared and subjected to a QTL-Seq approach (Takagi et al., 2013). This approach combines bulk-segregant analysis (Michelmore et al., 1991) with NGS technologies to locate candidate genomic regions more rapidly than traditional linkage mapping. A total of 34 F_2_ plants with red fruits and 32 F_2_ plants with white fruits were selected for bulking DNA in two pools referred as RF and WF, respectively. Next, we performed Illumina high-throughput sequencing of each pool to a 50× genome coverage and aligned short reads to the *F. vesca* reference genome v4.0.a1 (Edger et al., 2018). Frequencies for SNPs in each pool were calculated and average SNP-indexes were computed in 3-Mb intervals. The difference in SNP allele frequencies between the RF and WF pools (ΔSNP-index) were plotted against the *F. vesca* genome (Figure 1A). At the 97% confidence level, significant SNPs (in blue) were only detected on chromosome 1 at intervals 7,970,000-14,800,000 and 23,450,000-23,870,000 bp. We next extended the analysis to small and large INDELs in addition to SNP variants. Within significant intervals we searched for genes with non-synonymous variants and ΔSNP-indexes higher than 0.5 and homozygous (SNP-index = to 0 or 1) in the WF pool. The target intervals we identified harbored 105 candidate genes (Supplemental Table 1), including FvH4_1g22020 *(FvMYB10).* Bioinformatic analysis for large insertions detected a 52 bp insertion in *FvMYB10* in the WF pool in comparison to the RF pool and the reference *F. vesca* genome. Further analysis was focused on the 52 bp INDEL located at the beginning of the third exon of *FvMYB10*, the strongest candidate mutation underlying the white-fruited phenotype.

**Figure 1.**
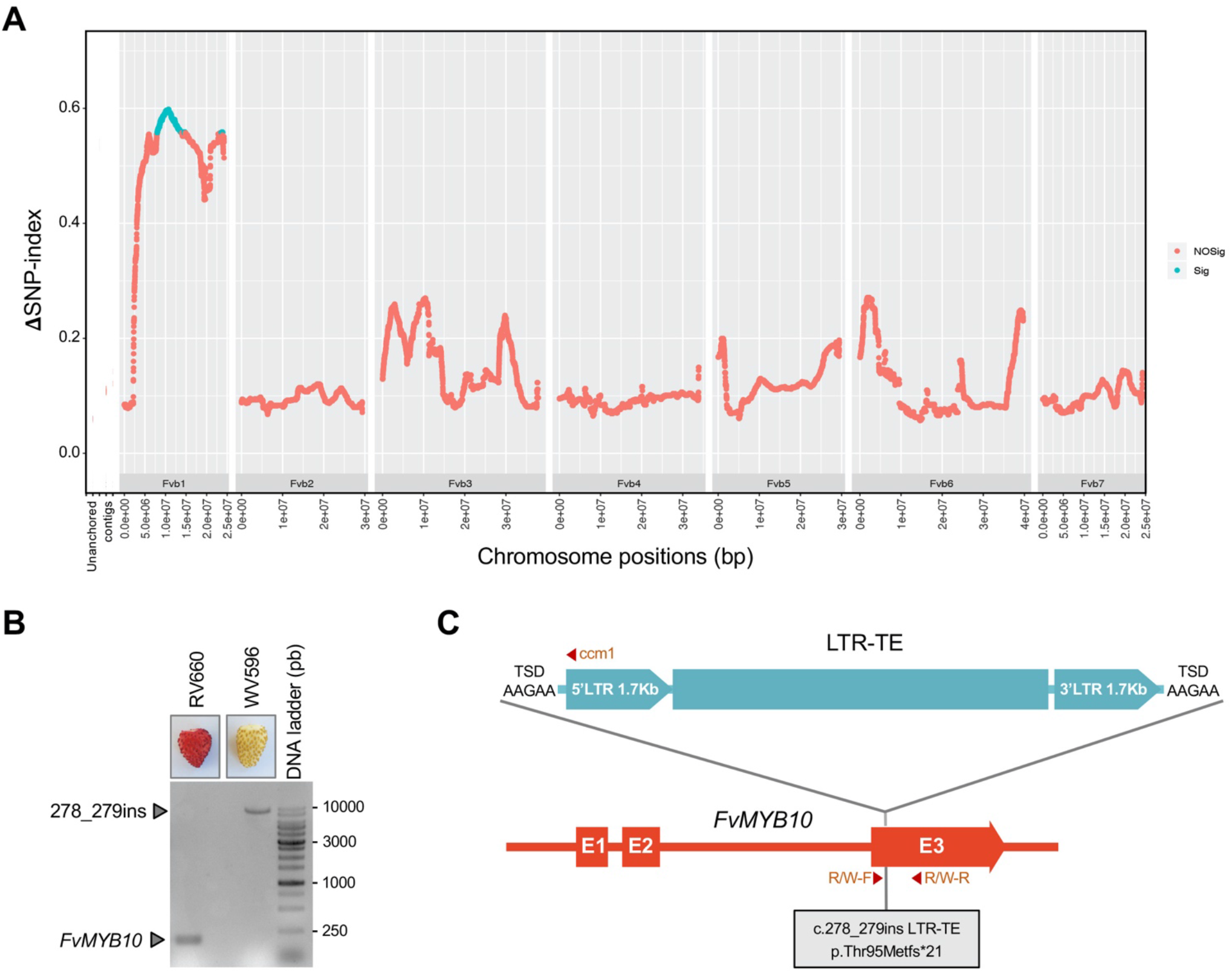
Quantitative trait loci (QTL)-Seq identifies a significant region on Fvb1 controlling fruit color in RV660 × WV596. (**A**) ΔSNP-index plot (Y-axis) over the seven *F. vesca* chromosomes. Line represents a sliding window of 3 Mb at 10 kb intervals. Significant SNPs are highlighted in blue (*p* < 0.03). (**B**) *FvMYB10* PCR amplification using primers R/W-F and R/W-R designed flanking the 52 bp INDEL (c.278_279ins). In RV660 the expected fragment size was 181 bp according to the reference ‘Hawaii-4’ genome. See scheme in (C). (**C**) Schematic representation of FvMYB10-*gypsy*, the Long Terminal Repeat Transposable Element (LTR-TE) identified in WV596 *FvMYB10* coding sequence. In the scheme, primers used for population genotyping have been highlighted (R/W-F, R/W-R and ccm1, the latter at the 5’ end of the LTR-TE; Supplemental Figure 2).

### *FvMYB10* coding region in white-fruited WV596 carries an LTR-TE insertion that impairs anthocyanin accumulation

Primers flanking the 52 bp insertion were designed and used to amplify the corresponding genomic fragment from white-fruited accession WV596 and the red-fruited cultivar ‘Reine des Vallées’ (RV660). Standard PCR generated the expected 181 bp product from RV660 but failed to generate a product in WV596. Reanalysis using the long-range 5-Prime PCR Extender System amplified a 10 kb product from WV596 (Figure 1B). Cloning and Sanger sequencing of the 10 kb genomic fragment from WV596 allowed us to confirm the insertion at position 278 of the *FvMYB10* cDNA but it was larger than initially predicted. Further sequencing by primer walking of the entire 10 kb fragment revealed the insertion located in the third exon of WV596 *FvMYB10* corresponds to a Long Terminal Repeat Transposable Element (LTR-TE), 9,509 bp in length, belonging to the Gypsy subfamily. This novel *FvMYB10* allele was designated *fvmyb10-2,* and the LTR-TE was named FvMYB10-*gypsy.* Upon insertion, FvMYB10-*gypsy* generated the 5 bp AAGAA target site duplication. Two almost identical (one nucleotide substitution) 1.7 kb Long Terminal Repeats (LTRs) were identified flanking the ORFs necessary for TE replication and transposition (Figure 1C). To determine how tightly linked *fvmyb10-2* was with the white fruit phenotype in the RV660 × WV596 F_2_ population, a reverse primer binding to FvMYB10-*gypsy* 5’ terminal repeat was used to genotype the population in combination with primers flanking the insertion point (Figure 1C). The *fvmyb10-2* allele co-segregated with the white phenotype in the entire population (Supplemental Figure 2) following the expected 1:2:1 ratio for a single-gene codominant trait.

*FvMYB10* transcript accumulation was not altered in the WF pool upstream of the insertion site but, as expected, it was abolished downstream of the insertion site (Figure 2A). In addition, the LTR-TE sequence introduced a series of premature stop codons in *FvMYB10* coding sequence. The predicted truncated protein lacks the C-terminal-conserved motif KPRPR[S/T]F for Arabidopsis anthocyanin-promoting MYBs (Stracke et al., 2001), also found in *Rosaceous MYB10* and known anthocyanin MYB regulators from other species (Lin-Wang et al., 2010). As a result, *fvmyb10-2* is expected to be non-functional and unable to induce the expression of FvMYB10 target genes from the anthocyanin pathway. Therefore, the expression of some representative FvMYB10 targets in the *fvmyb10-2* background was tested using qRT-PCR in ripe fruits from the RF and WF pools. As shown in Figure 2A, transcript accumulation from the downstream structural genes *CHI, F3H, DFR, ANS/LDOX* and *UFGT* were significantly reduced in ripe white fruit, confirming *fvmyb10-2* is a loss-of-function allele.

**Figure 2.**
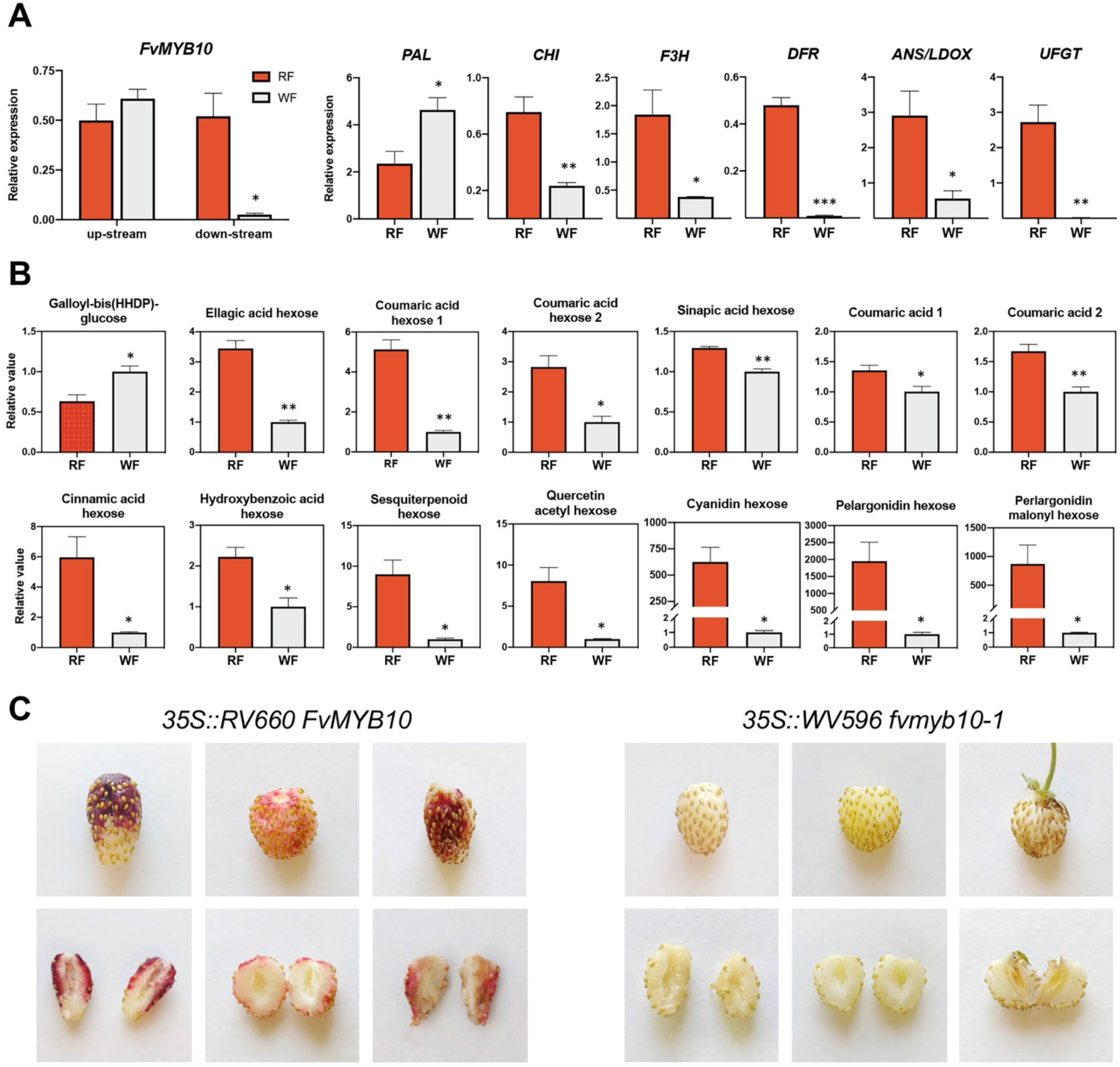
Effect of *FvMYB10* mutation on *F. vesca* structural genes and metabolites. **(A)** Expression by qRT-PCR of *FvMYB10* and anthocyanin structural genes in fruits from RF and WF F_2_ pools. For *FvMYB10,* primers were designed upstream (left) and downstream (right) LTR-TE insertion point (Supplemental table 9). Results are expressed as the average of three independent biological replicates. Error bars express standard error of the mean. Asterisks indicates significant differences as determined by Student’s *t* test (\**p*<0.05; \*\**p*<0.01; \*\*\**p*<0.005). **(B)** Secondary metabolites with significant differences between pools. Average values relative to WF pool. Asterisks indicates significant differences as determined by Student’s *t* test (\**p*<0.05; \*\**p*<0.01). **(C)** Transient RV660 *FvMYB10* overexpression restored anthocyanin biosynthesis in WV596 fruits. Three representative fruits are shown. As a control, fruits were agroinfiltrated with the truncated *fvmyb10-2* gene, which failed to induce anthocyanin accumulation.

To identify which secondary metabolites were affected by *FvMYB10* downregulation, we analyzed ripe fruits in the WF and RF pools by Ultra Performance Liquid Chromatography coupled to Tandem Mass Spectrometry (UPLC-Orbitrap-MS/MS). The use of the two bulked pools of F_2_ lines instead of parental lines RV660 and WV596, allowed identification of secondary metabolites affected only by *FvMYB10* down-regulation, excluding those metabolites differing in the two accessions due to variation in other genes. A total of 88 metabolites, including 30 ellagitannins, 41 flavonoids, 11 hydroxycinnamic acid derivatives, three hydroxybenzoic acid derivatives, and three terpenoids were identified (Supplemental Table 2). Significant differences were detected for only 14 metabolites, and the levels of all but one metabolite were increased by FvMYB10 (Figure 2B). As expected, cyanidin hexose and pelargonidin hexose, the two main anthocyanins, were 625- and 1,950-fold higher in the RF pool, respectively. Mutation of *FvMYB10* reduced the level of one terpenoid (sesquiterpenoid hexose) and one benzoic derivative (hydroxybenzoic acid-hexose) by 2.2- and 9-fold, respectively. Interestingly, the mutation in *FvMYB10* reduced ellagic acid hexose by 3.4-fold while galloyl-bis(HHDP)-glucose was increased 58%. Finally, one quercetin derivative and six hydroxycinnamic acid derivatives were decreased in the WF pool. Effects in the same direction on quercetin derivatives and coumaric acid derivatives were observed when *FaMYB10* was transiently down-regulated in octoploid strawberry (Medina-Puche et al., 2014) or when *FvMYB10* was engineered in *F. vesca* (Lin-Wang et al., 2014). Other studies have compared secondary metabolites of fruit from cultivars differing in fruit color and again quercetin derivatives, coumaric and cinnamic hexoses and ellagic acid were also associated with MYB10 function (Härtl et al., 2017; Wang et al., 2020). These results show that a very limited number of metabolites are affected by MYB10 regulation, suggesting a primary influence on anthocyanin biosynthesis and few effects in specific flavonols, ellagitanins and hydroxycinnamic acids. The reduction in these metabolites did not affect total antioxidant capacity of fruits (Supplemental Figure 1). Similarly, no significant differences were observed between RF and WF pools for total soluble solids content (SSC), titratable acidity or ascorbic acid content (Supplemental Figure 1), indicating that white fruits resulting from downregulation of *MYB10* are similarly rich sources of nutritional compounds other than anthocyanins.

Finally, we functionally validated *fvmyb10-2* as the causal agent of the lack of anthocyanin accumulation by transient overexpression in WV596 fruits. Two constructs were generated to test the complementation: *35S_pro_*:RV660 *FvMYB10* and *35S_pro_:WV596 fvmyb10-2.* Only RV660 *FvMYB10,* the allele from the red fruited ‘Reine des Vallées’ was able to induce anthocyanin accumulation in fruits, but not WV596 *fvmyb10-2* (Figure 2C), further confirming *fvmyb10-2* as the causal gene of the white fruit phenotype. Previous studies using different *F. vesca* accessions identified an independent polymorphism (G36C) in *FvMYB10* as the underlying cause of anthocyanin-less fruits (Zhang et al., 2015; Hawkins et al., 2016). This specific SNP is translated into a W12S amino acid substitution at a conserved residue within the R2 DNA-binding domain and has been identified in five white-fruited *F. vesca* accessions: Hawaii-4, ‘Yellow Wonder’, ‘Pineapple Crush’, ‘White Soul’ and ‘White Solemacher’. All of them have the C nucleotide and thus the W12S substitution in FvMYB10 (Hawkins et al., 2016). This polymorphism, named *fvmyb10-1* in this study, was not found in WV596, which had the wildtype G nucleotide.

### Independent *FvMYB10* mutations explain the lack of anthocyanins in diverse *F. vesca* ecotypes

To assess the incidence of *fvmyb10-1 or fvmyb10-2* alleles on fruit color natural variation in *F. vesca,* we broadened our analysis to include accessions with white fruits from different geographic origins: Hawaii-4, ‘Yellow Wonder’, ‘Pineapple Crush’, ‘White Soul’ and ‘White Solemacher’, GER1, GER2, ‘South Queen Ferry’, UK13, SE100 and FIN12 (Supplemental Table 3). Our PCR marker revealed that FvMYB10-*gypsy* insertion *(fvmyb10-2* allele) was not present in any of the accessions analyzed besides WV596 (Figure 3A). In FIN12, we could not detect the FvMYB10-*gypsy* associated band nor the endogenous *FvMYB10* gene (Supplemental Figure 3A), suggesting a putative large rearrangement in the surrounding chromosomal region.

**Figure 3.**
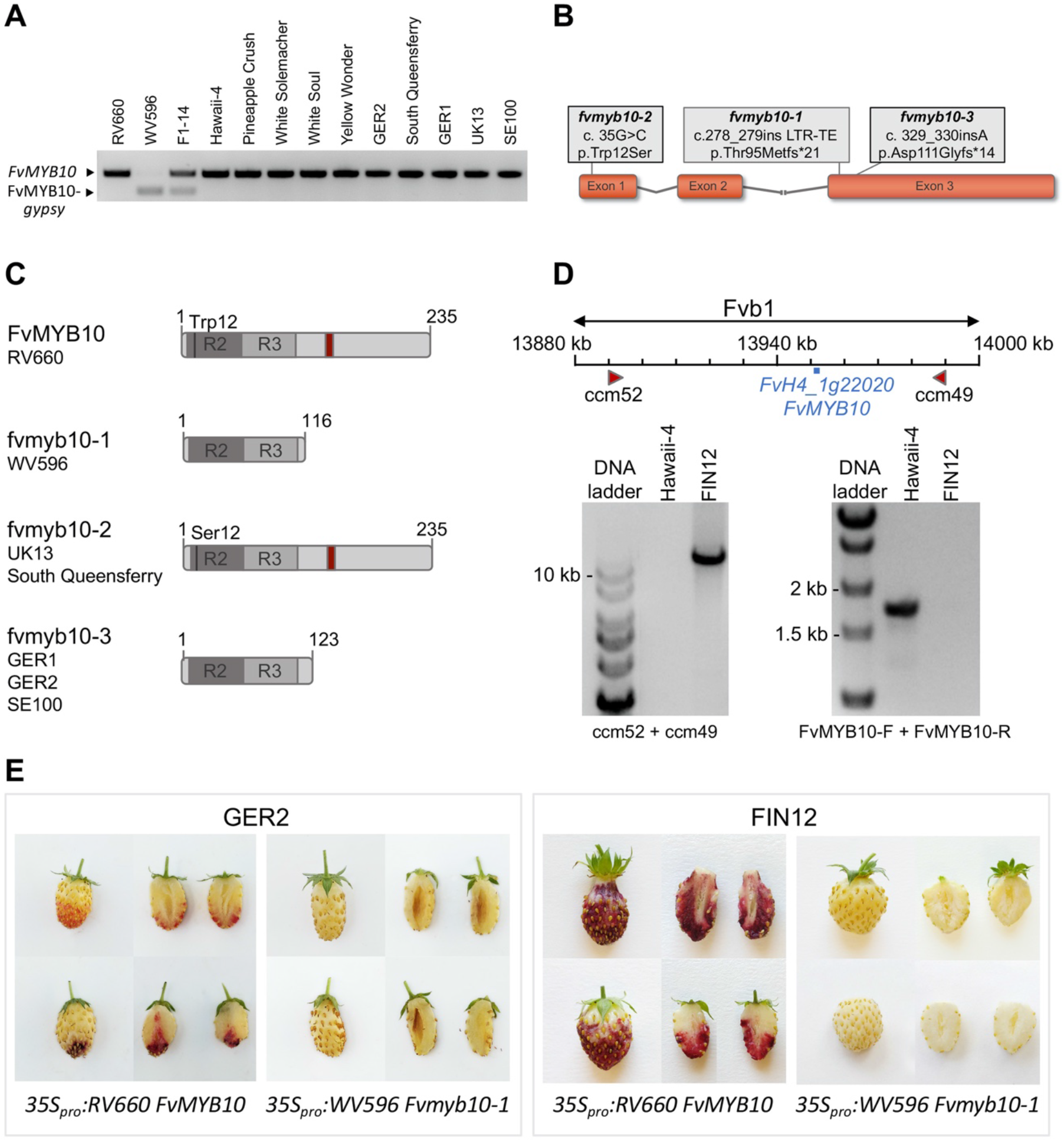
Multiple *FvMYB10* variants were detected in a panel of eleven white fruited *F. vesca* ecotypes. **(A)** FvMYB10-*gypsy* insertion *(fvmyb10-2)* was not detected in *FvMYB10* from other white fruited *F. vesca* ecotypes. **(B)** *FvMYB10* schematic representation showing the three alleles identified in white *F. vesca* ecotypes. *fvmyb10-2* and *fvmyb10-3* were identified in this study whereas *fvmyb10-1* has been previously described (Zhang et al., 2015; Hawkins et al., 2016). **(C)** Outline of the protein products encoded by *FvMYB10* alleles represented in (B) and list of the accessions in which they were found. **(D)** FIN12 has a large deletion in Fvb1 resulting in the loss of 7 predicted genes (Supplemental Figure 3). *FvMYB10* is among them and has been highlighted in the scheme. Primers ccm52 and ccm49 were designed flanking the region In Hawaii-4 both primers are 100 Kb apart, resulting in no PCR product. In FIN12 a >10 kb band was obtained. Right picture shows *FvMYB10* CDS amplification with primers FvMYB10-F and FvMYB10-R only in ‘Hawaii-4’ but not in FIN12. **(E)** *FvMYB10* transient overexpression assays showing GER2 and FIN12 white fruit complementation. Two representative fruits are shown.

White-fruited phenotypes in Hawaii-4, ‘Yellow Wonder’, ‘Pineapple Crush’, ‘White Soul’ and ‘White Solemacher’ are known to carry the *fvmyb10-1* allele (Hawkins et al., 2016). From the rest of the accessions, GER1, GER2, ‘South Queen Ferry’, UK13, SE100 and FIN12, *FvMYB10* CDS was cloned and Sanger sequenced. Sequence analysis revealed a third allelic variant previously not described, *fvmyb10-3,* in the accessions from Northern Europe GER1, GER2, and SE100. The *fvmyb10-3* allele had an A nucleotide insertion at position 329 of the cDNA (c. 329_330insA). The predicted protein contains the Asp111Gly substitution and a frameshift generating a premature stop codon 14 residues downstream from the insertion site (Figure 3B and C). Similar to *fvmyb10-2,* the resulting 123 aa protein transcribed from *fvmyb10-3* lacks the C-terminal domain found in anthocyanin related MYB factors (Stracke et al., 2001; Lin-Wang et al., 2010). Next, the British accessions ‘South Queen Ferry’ and UK13, both carried the previously published *fvmyb10-1* allele with the G36C SNP. In the accession FIN12 we were not able to amplify the *FvMYB10* coding region, but whole genome resequencing of FIN12 revealed an extensive region (~ 100 kb) in FIN12 chromosome one (Fvb1) with extremely low sequence coverage (Supplemental Figure 3B) suggesting a large deletion affecting this chromosomal fragment. The uncovered region of FIN12 corresponds approximately to positions 13,890,000-13,990,000 bp from the reference Hawaii-4 genome which contains seven predicted coding sequences, including *FvMYB10* (Figure 3D). Primers designed in the predicted flanking regions detected a ~10kb band in FIN12 but none in Hawaii-4 (Figure 3D), confirming the large deletion from FIN12 Fvb1.

To demonstrate that (1) *fvmyb10-3* in GER1, GER2 and SE100 and (2) the FIN12 large deletion from Fvb1 were the causal mutations leading to white fruits, we tested whether a functional *FvMYB10* copy was able to restore fruit pigmentation when transiently expressed in fruits from those accessions. The same constructs described for *fvmyb10-2* complementation in WV596 were employed in this assay. Once again, expression of full-length RV660 *FvMYB10* allele was sufficient to induce anthocyanin accumulation in all white-fruited accessions (Figure 3E). Interestingly, anthocyanin accumulation was observed in the fruit epidermis as well as in the inner receptacle, where red color is not observed in wild type fruits. This phenomenon was also observed in WV596 in our study and by Hawkins et al. (2016) when complementing ‘Yellow Wonder’ fruits. This probably resulted from the use of a constitutive 35S promoter, suggesting that *FvMYB10* might not normally be expressed in the internal tissues.

In summary, we have identified three *FvMYB10* mutations, in addition to the previously described *fvmyb10-1* allele (Hawkins et al., 2016), which explain the lack of pigmentation in the skin of several white-fruited *F. vesca* accessions: (1) An LTR-TE insertion at the third exon of *FvMYB10 (fvmyb10-2);* (2) a single nucleotide (A) insertion at position 329 of *FvMYB10* cDNA *(fvmyb10-3);* and (3) a large deletion in chromosome 1 that removed seven predicted genes, including *FvMYB10.* The different allelic variants affecting fruit color here described are summarized in Supplemental Table 3. Notably, none of the white-fruited *F. vesca* accessions we studied carried a functional *FvMYB10* gene. This indicates that the white-fruited phenotype in *F. vesca* arose through different independent mutations in the same *MYB* gene, illustrating a convergent/parallel evolutionary mechanism. White-fruited strawberry mutants targeting other genes but *MYB10* has been artificially obtained, as the *reduced anthocyanins in petioles (rap)* mutant, which was identified in a mutagenized *F. vesca* population (Luo et al., 2018). The *RAP* gene encodes a glutathione S-transferase (GST) that binds anthocyanins to facilitate their transport from the cytosol to the vacuole. Similar white fruit and white stem phenotypes were observed in cultivated strawberry after *RAP* knockout using CRISPR/Cas9 (Gao et al., 2020). The fact that *rap* mutants, nor others in different genes, have not been identified in nature led us to speculate that mutations in genes that result in a general lack of anthocyanins are negatively selected due to its role in protecting plants against a wide range of abiotic stresses (Tohge et al., 2017), while lack of anthocyanins only in fruits might not be as detrimental.

### Detection of QTLs controlling fruit color in octoploid strawberry

Improvement of cultivated strawberry is more challenging than in the diploid *F. vesca,* not only because of its octoploid genome but also because frequent homoeologous exchanges have occurred following polyploidization, replacing substantial portions of some subgenomes with sequences derived from ancestrally related chromosomes (Edger et al., 2019a). These exchanges have been found to be biased towards the *F. vesca*-like subgenome, although they are not completely unidirectional (Edger et al., 2019a). It is therefore crucial to characterize the genetic control of color variation within the octoploid *Fragaria* species, and to answer questions such as: how many genes are involved in controlling this trait, are all possible homoeologous copies present in the genome, are they being expressed, and are previously identified mutations in *FvMYB10* also present in octoploid strawberry?

To accomplish this objective, we studied two diverse octoploid populations characterized by a broad variation in fruit skin (University of Florida strawberry breeding population 17.66) and flesh color (Hansabred SS×FcL population). The 17.66 population is an F_1_ derived from biparental cross between two UF advanced selections (FL 13.65-160 and FL 14.29-1) that segregates for white skin color (Supplemental Figure 4). Within this population, seedlings with white or light pink skin are also characterized by white internal flesh. Using the second-generation high-density 50K SNP Array (FanaSNP) (Hardigan et al., 2020), a genome-wide association study (GWAS) was conducted to identify the chromosomal regions and specific SNPs associated with skin color in this breeding population. Association analyses of 43,422 SNP markers with 95 genotypes were performed to detect marker-trait associations with three different analysis models (GLM, LMM, and MLMM), and SNPs strongly associated with white fruit skin color were located on chromosome Fvb1-2 (Figure 4A). The most significant SNP marker was probe AX-184080167 in all three models and was adjacent to a MYB10 homoeolog (Fvb1-2 cv. Camarosa, 15,395,876 bp).

**Figure 4.**
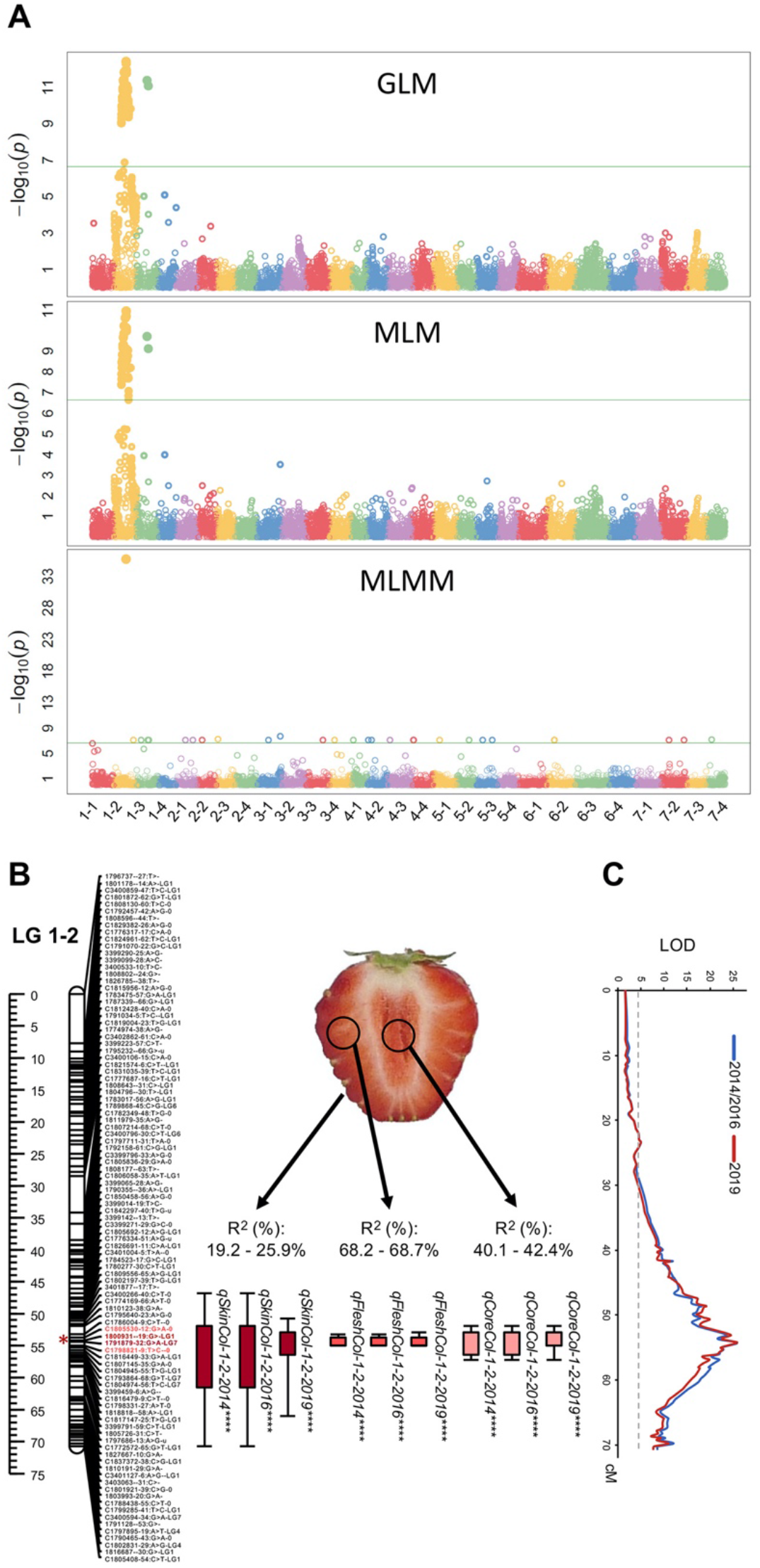
SNPs and QTLs associated with fruit color traits in octoploid strawberry. **(A)** Manhattan plots of a general linear model (GLM), a mixed linear model (MLM), and a multiple-locus mixed model (MLMM) for white fruit in UF breeding population 17.66. The different colors indicate plots for different chromosomes, which follow the order: chromosome 1-1 to chromosome 7-4. Green line represents the significance threshold. **(B)** Positions of QTLs on LG 1-2 controlling fruit color detected in the ‘Senga Sengana’ × *F. chiloensis ssp. lucida* USA2 F_2_ population. Thick and thin bars represent 1- and 2-LOD QTL intervals, respectively, and are drawn at the right of linkage group 1-2. Names of QTL as described in Supplemental table 4. Markers in the pick and 2-LOD interval are highlighted in red and pink, respectively. Estimated position of *FvMYB10* is indicated with an asterisk based on the position of flanking SNPs in *F. vesca* reference genome v.4. **(C)** LOD scores of flesh QTL are plotted along the x-axis (dotted line indicates the 4.5 LOD threshold), whilst genetic distances are plotted along the y-axis.

The second population used in this study, SS×FcL, is an F_2_ derived from the interspecific cross between *F. ×ananassa* ‘Senga Sengana’ and *F. chiloensis* ssp. *lucida* USA2 that segregates for skin and flesh color in the fruits. While all 105 F_2_ individuals accumulated varied levels of anthocyanins in the skin, the variation in fruit flesh color was somewhat qualitative, with a number of F_2_ individuals displaying white flesh (Supplemental Figure 5). The population was phenotyped for skin and flesh color over three seasons and genotyped using DArTseq markers previously developed for octoploid strawberry (Sánchez-Sevilla et al., 2015). A linkage map comprising 2,991 SNPs and covering a total length of 2,377.21 cM was generated (Supplemental Figure 6). Three QTLs *(qSkinCol-1-2, qSkinCol-3-1* and *qSkinCol-2-3)* were detected for fruit skin color. The first QTL was detected in all three years, while the other two were only detected in one or two years (Supplemental Table 4). The phenotypic variance contributed by each QTL ranged from 12.2 to 25.9%. Previous studies have detected QTLs for color traits in LGs belonging to Homoeology group (HG) 1 (Lerceteau-Kohler et al., 2012), 2 (Lerceteau-Kohler et al., 2012; Castro and Lewers, 2016; Zorrilla-Fontanesi et al., 2011), 3 (Lerceteau-Kohler et al., 2012; Zorrilla-Fontanesi et al., 2011), 5 (Zorrilla-Fontanesi et al., 2011; Castro and Lewers, 2016) and 6 (Castro and Lewers, 2016; Lerceteau-Kohler et al., 2012) although the different types of markers used in each study makes it difficult to compare LGs and positions. Interestingly, a major QTL for flesh color, *qFleshCol-1-2,* was detected in the three years in the SS×FcL population on LG 1-2 (Figure 4B, C). This QTL explained a high proportion of the phenotypic variation (68.2-68.7 %) and colocalized with *qCoreCol-1-2,* a QTL controlling 40.1-42.4 % variation of fruit core color (Figure 4B; Supplemental table 4). We concluded that the same gene was probably affecting fruit flesh and core color (internal color), and to a lesser extent, skin color in octoploid strawberry. The 2-LOD confidence interval of *qFleshCol-1-2* spanned a region on LG 1-2 from 53.2 to 55.0 cM (Supplemental Table 4) that corresponded to the region from 13.75 to 15.35 Mb on *F. vesca* chromosome 1 (v4.0.a1; (Edger et al., 2018). The 1.6 Mb internal color QTL confidence interval region was found to contain 171 annotated genes. Among them, once again *MYB10* was the most likely candidate as the gene underlying the QTL.

### *FaMYB10-2* is the dominant homoeolog in octoploid strawberry

As both the UF population 17.66 and SS×FcL studies identified *MYB10* from chromosome Fvb1-2 as a putative causal locus, we further investigated this homoeolog within an octoploid reference genome sequence. The *F. ×ananassa* ‘Camarosa’ reference genome (Edger et al., 2019a) contains four *FaMYB10* homoeologs. Chromosomes Fvb1-1 and Fvb1-2 carry one *FaMYB10* homoeolog each: *FaMYB10-1* (maker-Fvb1-1-snap-gene-139.18 or FxaC_4g15020) and *FaMYB10-2* (maker-Fvb1-2-snap-gene-157.15 or FxaC_2g30690), respectively. Two *FaMYB10* genes were found on chromosome Fvb1-3, which were designated *FaMYB10-3A* (maker-Fvb1-3-augustus-gene-143.29 or FxaC_3g25620) and *FaMYB10-3B* (maker-Fvb1-3-augustus-gene-144.30 or FxaC_3g25830), but only *FaMYB10-3B* has a full-length ORF*. FaMYB10-3A* cannot be functional in activating anthocyanin biosynthesis, at least in ‘Camarosa’, as *MYB10-3A* CDS is interrupted by a *Ty1-copia* retrotransposon insertion at the end of the second intron. As a result, a truncated MYB10 protein lacking the 152 C-terminal residues is predicted. Lastly, no *FaMYB10* allele was found on chromosome Fvb1-4. Therefore, besides *FaMYB10-2,* located at the same position as identified QTL in both octoploid populations, other *MYB10* copies could potentially be functional, as they encode full-length MYB proteins. However, alignment of transcriptomic sequences from a previous RNA-seq study (Sánchez-Sevilla et al., 2017) to the chromosome-scale ‘Camarosa’ genome (Edger et al., 2019a) allowed us to obtain sub-genome specific global expression profiles. This sub-genome specific expression analysis revealed that *FaMYB10-2,* in the *F. iinumae-derived* subgenome, is the dominant homoeolog throughout fruit development in strawberry receptacle and achene tissues (Supplemental Figure 7). In turning and red receptacles, for example, where *FaMYB10* expression peaks, the expression of *FaMYB10-2* represents 97% of total *FaMYB10* expression in the respective ripening stage. By contrast, transcript accumulation from the other two full-length homoeologs from chromosomes Fvb1-1 and Fvb1-3, *MYB10-1* and *MYB10-3B*, was barely detectable, accounting for only 0.6% of total *FaMYB10* expression (Supplemental Figure 7). Similarly, re-examination of *MYB10* expression from a previous study using 61 strawberry lines also demonstrates that *FaMYB10-2* is the dominantly expressed homoeolog compared to those on Fvb1-1 and Fvb1-3 (Barbey et al., 2019). Thus, we expected that a non-functional FaMYB10-2 would be enough to abolish anthocyanin pathway induction in octoploid strawberry.

### *MYB10* allelic variants in white-fruited octoploid accessions

To identify sequence variation in *MYB10* gene between white- and red-skinned fruits from the UF breeding population 17.66, we performed NGS-based bulked-segregant analysis (BSA). *FaMYB10* cDNAs amplified from pools of white- and red-fruited accessions were sequenced using Illumina MiSeq platform. A total of 242,900 and 236,178 reads were generated in the white and red fruit pools, from which 93.85% and 88.24% were mapped in the *FaMYB10-2* gene. In white fruit accessions we found an 8 bp insertion (ACTTATAC) at position 491 of the *FaMYB10-2* ORF generating a premature stop codon and a predicted 179 amino acid protein instead of the 233 amino acid wild type FaMYB10-2 (Supplemental Figure 8). Deletion of the 54 C-terminal residues may render FaMYB10 inactive. In fact, this same polymorphism was recently shown to be associated with white fruits in ‘Snow Princess’, and restoration of anthocyanin biosynthesis was only possible by over-expression of WT *MYB10* (Wang et al., 2020). The presence of the same mutation, hereafter named *famyb10-1,* suggests that the white-fruited selection FL 14.29-1 and ‘Snow Princess’ may share a common pedigree.

Unlike in *F. vesca* accessions or in *F. ×ananassa* line FL 14.29-1, *MYB10* coding sequence comparison between ‘Senga Sengana’ and USA2, the parents of the SS×FcL F_2_ population, revealed no polymorphisms potentially affecting protein stability/functionality. In fact, anthocyanins are accumulated in fruit skin in all F_2_ individuals suggesting the expression of a functional MYB10 gene in this tissue. In order to examine the presence of structural variation in ‘Senga Sengana’ and USA2 *MYB10* promoters *(MYB10_pro_)*, primers were designed based on the ‘Hawaii-4’ *F. vesca* reference genome (Shulaev et al., 2011) intended to amplify a ~1 kb product, from −941 to +55 relative to the *FvMYB10* translational start codon (Figure 5A). PCR was performed in two red-(‘Camarosa’ and ‘Senga Sengana’) and 3 white-fleshed genotypes (USA1, USA2 and FC157). All the accessions tested showed the expected ~1 kb amplicon together with an unforeseen band of 1.6 kb. In accessions with white-fleshed fruits an additional extra band of 2.1 kb in USA1 and USA2 or 2.8 kb in FC157 emerged (Figure 5B). PCR products from ‘Senga Sengana’, USA2 and FC157 were cloned and sequenced, confirming they all were *MYB10_pro_* alleles. In order to assign each *MYB10_pro_* variant to its respective subgenome, they were aligned with the upstream regulatory sequences of the four ‘Camarosa’ *FaMYB10* homoeologs. Binding sequences of the primers used for *MYB10_pro_* amplification were localized in all four ‘Camarosa’ homoeologs and the sequence comprised between them was retrieved and used for alignment and homology tree construction together with the different sequenced alleles from ‘Senga Sengana’, USA2 and FC157 (Figure 5C). The alignment revealed that the region upstream of *MYB10* is extremely polymorphic, containing a high density of SNPs, INDELs and transposon-derived sequences. The initially expected ~1 kb product corresponds to the 938 bp long *FaMYB10-3A_pro_*. All *MYB10-3A_pro_* alleles from the different backgrounds were grouped in the same clade. All of them were very similar in length (924-938 bp) and presented a high degree of sequence conservation (97-99% identity). The 1.6 kb allele was common to *MYB10-1,* in chromosome Fvb1-1, and *MYB10-3B* from Fvb1-3. Both *MYB10-1_pro_* and *MYB10-3B_pro_* from ‘Camarosa’ were identical and shared a 90% identity in the overlapping region with *MYB10-3A_pro_,* however *MYB10-3B_pro_* and *MYB10-1_pro_* had a 710 bp insertion at position −276 from the initial ATG (green segment in Figure 5C). Interestingly, *MYB10-3A_pro_* shows higher homology with *F. vesca FvMYB10_pro_* than to *MYB10-3B_pro_*. This relationship might reflect an event of chromosome translocation from Fvb1-4, which did not retain the copy of *MYB10,* rather than a duplication of *MYB10-3B* (or *MYB10-3A)* as, according to (Edger et al., 2019a), the octoploid chromosome Fvb1-4 originated from the diploid *F. vesca* progenitor. As observed for *MYB10-3A_pro_, MYB10-1_pro_* and *MYB10-3B_pro_* alleles from the different accessions were almost identical in length (1641-1649 bp) and sequence (96-97% identity) and grouped together in the same clade of the homology tree. Lastly, the fourth *FaMYB10_pro_* allele from the reference ‘Camarosa’ Fvb1-2 chromosome, *FaMYB10-2_pro_,* turned out to be much longer than the other three homoeologs, being almost 23 kb long. This size prevented *FaMYB10-2_pro_* PCR amplification from ‘Senga Sengana’ and ‘Camarosa’ under routine PCR conditions. Nonetheless, we were able to amplify *MYB10-2_pro_* alleles from white fleshed accessions, which were significantly shorter than 23 kb: 2.1 kb in USA1 and USA2, and 2.8 kb in FC157. USA2 *MYB10-2_pro_* was highly similar to USA2 *MYB10-1_pro_* and *MYB10-3B_pro_* 1.6 kb alleles (94% identity) but it had a tandem duplication of a 471 bp sequence. The first unit of the tandem duplication was a *MYB10-1_pro_* / *MYB10-3B_pro_* specific sequence (represented in light green in Figure 5C) and the second unit is colored in blue in the same scheme, highlighting that it is Fvb1-2-specific. On the other hand, FC157 *MYB10-2_pro_* was almost identical (98% identity) to USA2 *MYB10-2_pro_*, but FC157 had an additional 660 bp insertion disrupting one unit of the tandem duplicated sequence (dark blue segment in Figure 5C). Despite tremendous size differences, all *MYB10-2_pro_* variants were clustered in the same tree branch, as happened with the other *MYB10_pro_* homoeologs, and thus we can conclude that *MYB10_pro_* alleles from the same homoeolog from different accessions are phylogenetically closer to each other than the four alleles of a given background. It is noteworthy that *MYB10-2_pro_* is the allele that shows higher polymorphism among the different accessions, and the shorter *MYB10-2_pro_* alleles found in white-fleshed accessions are strong candidates to be the underlying polymorphisms at the *qFleshCol-1-2* QTL and the cause for white flesh in strawberry fruits.

**Figure 5.**
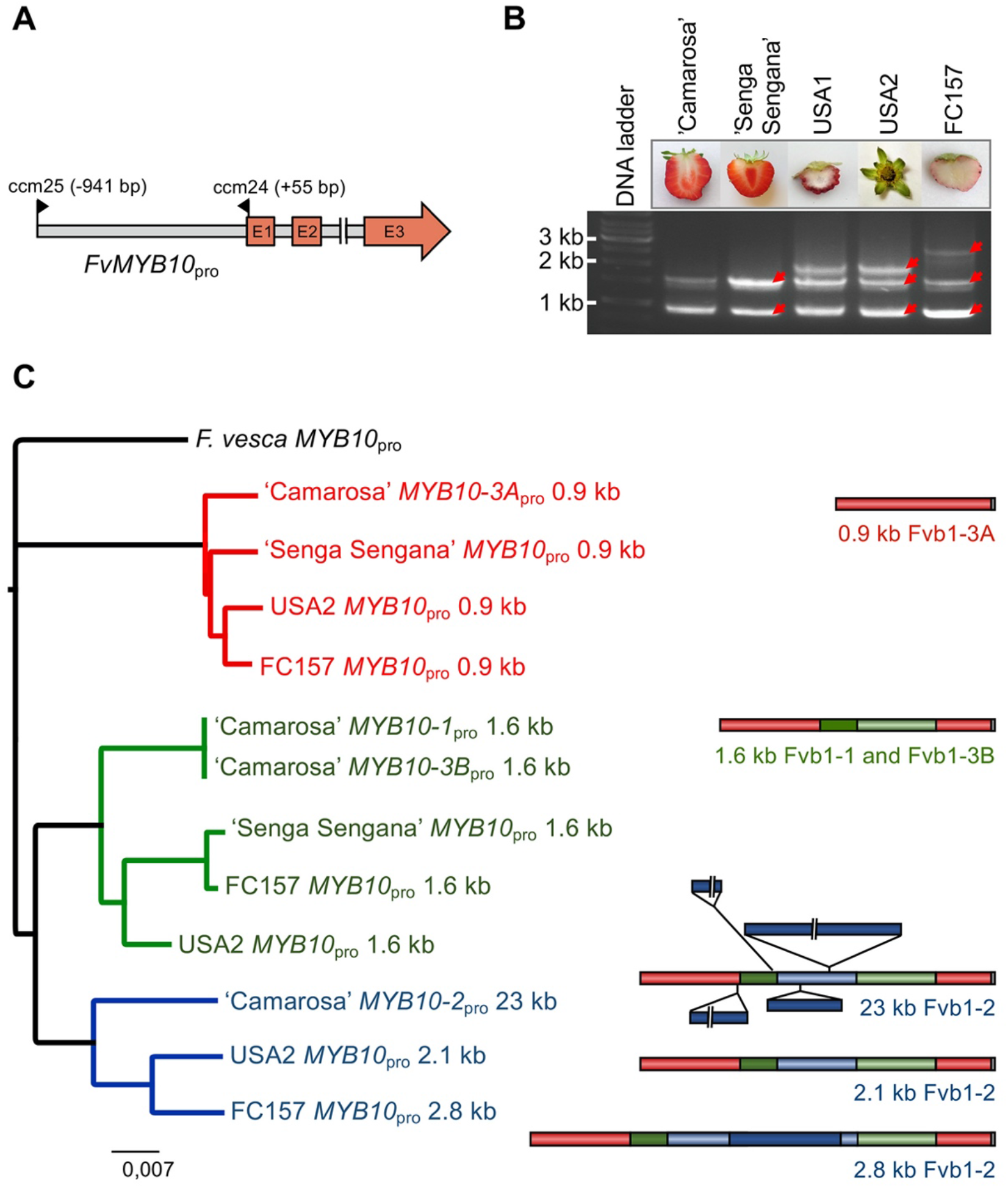
*MYB10* promoter variants across subgenomes and accessions in octoploid *Fragaria* spp. **(A)** *FvMYB10* schematic representation showing the position of primers used for *MYB10_pro_* PCR amplification from octoploid *Fragaria* spp. (Supplemental table 9). **(B)** *MYB10_pro_* PCR amplification yielded different sized promoter alleles from each accession. White-fleshed accessions USA1, USA2 and FC157 contained an additional 2.1- or 2.8-kb band compared to red-fleshed ones. Red arrows point to bands which were excised from the gel and cloned for sequencing. **(C)** Sequences obtained from (B) were aligned with the respective sequences of the four *FaMYB10* homoeologs from ‘Camarosa’ retrieved from the reference genome and used for homology tree construction. Sequences were grouped in three main clades which revealed their chromosomal origin: (1) Red clade contains promoter sequences from all Fvb1-3A homoeologs, (2) green clade from Fvb1-3B and Fvb1-1 homoeologs, which are identical, and (3) in blue, all alleles from Fvb1-2. On the right, a schematic representation of a promoter allele representative of each clade is shown. Three different versions of Fvb1-2 *MYB10_pro_* were found: (1) a 23 kb allele from ‘Camarosa’ containing four large INDELs, (2) a 2.1 kb allele from USA2 and (3) a 2.8 kb allele from FC157. A color code has been used for each homoeolog copy of *MYB10_pro_* in the tree and their respective hallmark sequences in schemes: Red for *MYB10-3A_pro_,* green for *MYB10-1_pro_* and *MYB10-3B_pro_* (they are identical) and blue for *MYB10-2_pro_*.

### A large transposon insertion at *MYB10-2* promoter is associated with the red-flesh phenotype

Sequence comparison among the 23 kb ‘Camarosa’ *MYB10-2_pro_* sequence and the corresponding alleles from the white fleshed accessions USA2 and FC157 revealed that the common region of ‘Camarosa’ *MYB10-2_pro_* shares 96% identity with alleles from both white-fleshed accessions. The substantial size differences were due to the presence of four large INDELs of 4,797 bp, 1,496 bp, 454 bp and 14,064 bp (Figure 5C). They are located 17.7 kb, 16 kb, 15,4 kb and 986 bp upstream of the ATG initiation codon of *MYB10-2*, respectively. Using Repbase and the software tool Censor (Kohany et al., 2006), the 4,797 bp and 14,064 bp insertions were identified as class II (DNA-type) TEs belonging to the CACTA family based on sequence similarity with the *F. vesca EnSpm-1_FV* element (Jurka, 2013; Shulaev et al., 2011). We designated them *FaEnSpm-1* (4,797 bp) and *FaEnSpm-2* (14,064 bp). Both elements are bordered by almost identical 14 bp terminal inverted repeat (TIR) sequences 5’-CACTACCAGAAAAT-3’, and several subterminal direct and inverted repeats were also identified. Upon insertion, *FaEnSpm-1* generated the 3-bp target site duplication (TSD) TCG, whereas *FaEnSpm-2* is flanked by the TSD CAA. *FaEnSpm-2* and *FaEnSpm-1* share important sequence similarity in their internal regions, but *FaEnSpm-1* is thought to be a defective deletion derivative as most of the sequences have been lost, including two transposase_21 domains (pfam02992) (Marchler-Bauer et al., 2015).

The presence of *FaEnSpm-2* at close proximity (< 1 kb away) to the *MYB10-2* coding region only in the red-fleshed accessions ‘Camarosa’ and ‘Senga Sengana’ prompted us to examine whether its loss correlates with the lack of anthocyanin accumulation in fruit flesh in the SS×FcL mapping population. A codominant Fvb1-2-specific PCR marker intended to be predictive for Internal Fruit Color (IFC-1 marker) was developed using a combination of three primers depicted in Figure 6A. When assayed in the entire population, it was able to predict the phenotype in >95% of the F_2_ individuals (Figure 6A). Depending on the season, the phenotypic score of 4-5 out of 105 F_2_ individuals did not match with their genotype. That subtle deviation might be explained by the observed variability among fruits from the same line, which could lead to misscoring of the phenotype. Alternatively, due to the quantitative nature of this trait, those individuals might represent the natural genetic variation not associated with the large-effect *qFleshCol-1-2* QTL.

**Figure 6.**
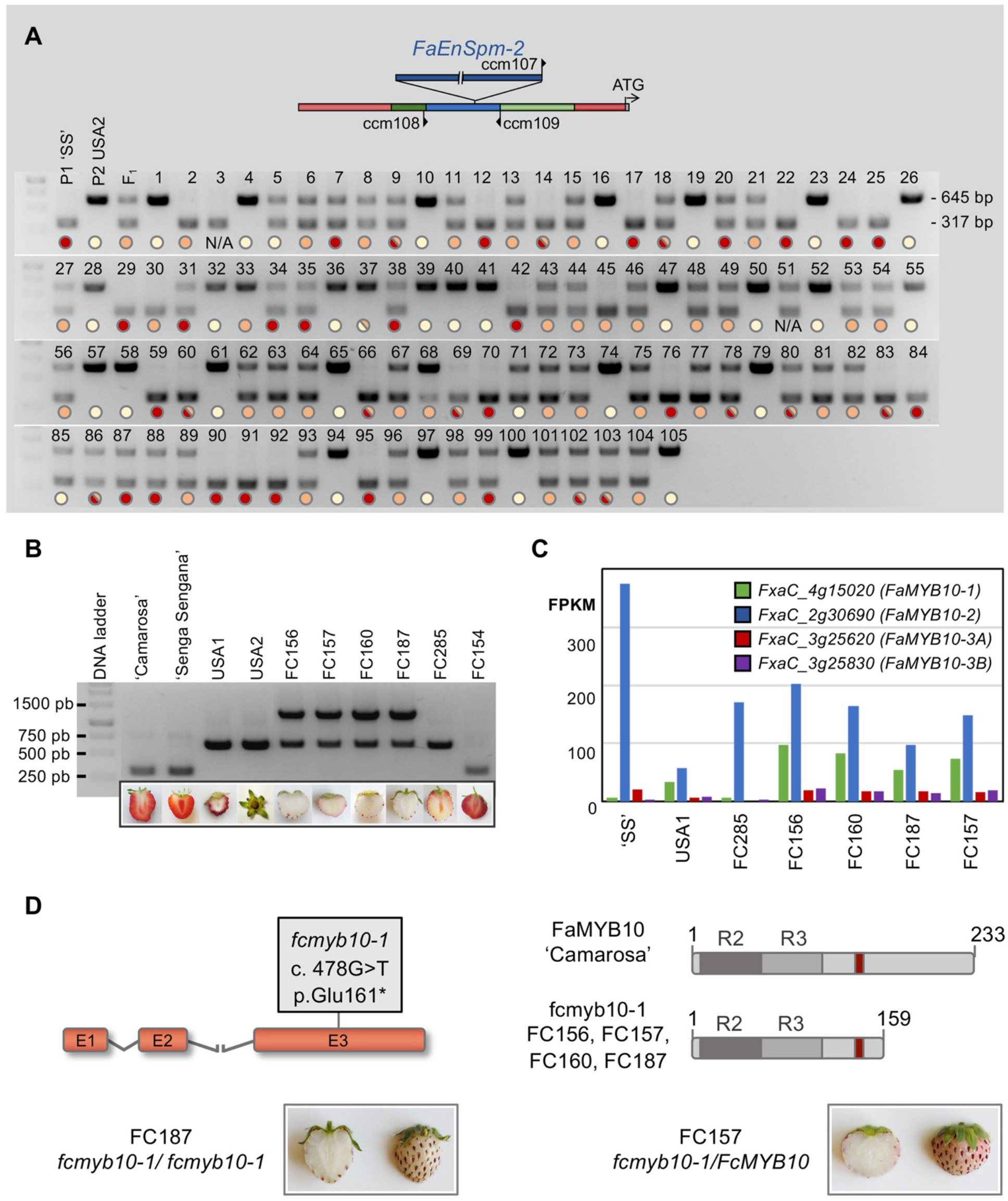
*FaEnSpm-2* insertion at *MYB10-2* upstream regulatory region is associated with internal red fruit color. Different mutations in the coding sequence lead to complete white fruits, lacking anthocyanins in the epidermis. **(A)** *FaEnSpm-2* co-segregates with red flesh color in the SS×FcL F_2_ population. The Fvb1-2-specific codominant IFC-1 marker was developed using a combination of three primers. Primer ccm107 binds to the junction sequence of *FaEnSpm-2* and together with ccm109 amplify a 317 bp band associated with red-flesh phenotype, whereas the 645 bp product (no TE insertion) associates with white flesh phenotype. Colored circles indicate flesh color: cream color=1, orange=2 and red=3. Circles with two colors represent different scores from different seasons (Supplemental Figure 5). **(B)** IFC-1 marker in *F. chiloensis* accessions. Only the red fleshed FC154 carried the *FaEnSpm-2* TE at *pMYB10-2.* Among white-fleshed *F. chiloensis* two different genotypes were found: (1) *F. chiloensis* ssp. *lucida* (USA1 and USA2) and *pacifica* (FC285) accessions both contain the smaller promoter allele version corresponding to the 2.1 kb *pMYB10* variant cloned from USA2, whereas *F. chiloensis* ssp. *chiloensis* accessions (FC156, FC157, FC160 and FC187 harbor the bigger 2.8 kb promoter allele identified in FC157. **(C)** Data from RNA-seq showing expression of each *MYB10* homoeolog in ‘Senga Sengana’ and six white-fleshed *F. chiloensis* accessions. Expression level dominance is biased towards *MYB10-2* in all accessions tested. **(D)** Scheme representing *fcmyb10-1* allele, which contains a G/T SNP in the coding sequence leading to a predicted truncated protein. Accessions homozygous for this variant in Fvb1-2, i.e. FC187, are completely white. FC157 is heterozygous *fcmyb10-1/FcMYB10* in chromosome Fvb1-2 (Supplemental Table 5) and has light pink epidermis.

The applicability of a given marker in breeding programs depends on its significance beyond a single cross. Therefore, the relevance of our IFC-1 marker and its ability to predict internal fruit color was tested in a wider set of white-fruited *F. chiloensis* accessions. To maximize genetic diversity, we selected accessions from ssp. *chiloensis* (FC156, FC157, FC160 and FC187), ssp. *lucida* (USA1), and ssp. *pacifica* (FC285; Supplemental Table 3; Figure 6B). Notably, the 317 bp band associated with the presence of *FaEnSpm-2* was also present in the *F. chiloensis* accession FC154 with red flesh but was absent from all white-fleshed accessions (Figure 6B). Furthermore, the marker allowed us to identify two different groups in the white-fleshed accessions, which also reflect their taxonomic relationships. All *F. chiloensis* ssp. *chiloensis* carried the 2.8 kb *MYB10_pro_* allele (1303 bp band) described for FC157. On the other hand, accessions from ssp. *pacifica* and *lucida,* taxonomically closer between them than to ssp. *chiloensis* (Staudt, 1999), carry the 2.1 kb *MYB10_pro_* allele (645 bp band).

### Expression of *MYB10* and anthocyanin biosynthetic genes are upregulated in red-fleshed accessions

TEs have the potential to alter or regulate the expression of proximal genes through multiple mechanisms including disruption of promoter sequences, introduction of novel alternative promoter sequences or epigenetic silencing (Vicient and Casacuberta, 2017; Hirsch and Springer, 2017; Rebollo et al., 2012). There are multiple examples where recruitment of a TE in the promoter region of *MYB10* orthologs leads to up-regulation of *MYB* expression and anthocyanin accumulation in other species such as orange, apple, pepper or brassica (Butelli et al., 2012; Jung et al., 2019; Yan et al., 2019; Zhang et al., 2019). Remarkably, in *B. oleracea* and pepper, the TE identified to enhance the expression of the host *MYB* gene is, as *FaEnSpm-2,* a CACTA element (Yan et al., 2019; Jung et al., 2019). To investigate the association of *FaEnSpm-2* with red flesh color, we profiled global gene expression using RNA-seq in fruits from ‘Senga Sengana’ and the six white-fleshed accessions previously genotyped in this study except for USA2. USA2 is a male plant and does not set fruits. Instead, we took advantage of its female sister line USA1, which presents the same phenotype segregating in the SS×FcL F_2_ population (Supplemental Table 3). Fragments per kilobase of transcript per million fragments mapped (FPKM) values for each *MYB10* homoeolog are presented in Figure 6C. As shown for ‘Camarosa’ (Supplemental Figure 7), in ‘Senga Sengana’ there is an obvious expression level dominance biased towards *MYB10-2.* Interestingly, *MYB10-2* expression was upregulated in ‘Senga Sengana’ compared to all white fleshed accessions. Total *MYB10* expression was lower in all white fleshed accessions too, although transcript accumulation from *MYB10-1* was higher in the four *F. chiloensis* ssp. *chiloensis* compared to ‘Senga Sengana’, USA1 or FC285, suggesting a putative mechanism of transcriptional compensation.

Comparative analysis of RNA-seq reads from the four *MYB10* homoeologs allowed us to identify SNPs and INDELs among all transcripts relative to the reference ‘Camarosa’ sequence (Supplemental Table 5). A total of 25 polymorphic nucleotides were identified in *MYB10* coding regions from the seven samples analyzed. Among them, 7 were silent mutations, 16 missense mutations at nonconserved residues and one nonsense mutation. None of the previously described polymorphisms in *FvMYB10* or the *FaMYB10* ACTTATAC insertion were present in the analyzed octoploid accessions. The nonsense mutation was only identified in *F. chiloensis* ssp. *chiloensis* accessions (FC156, FC157, FC160 and FC187) and consisted of a G to T transversion at position 478 of *MYB10* ORF. The substitution predicts a premature stop codon at amino acid position 160, generating a truncated protein lacking 74 residues from the C-terminal end (Figure 6C). This new *MYB10* allele derived from *F. chiloensis* was designated *fcmyb10-1.* It was found in the three full length *MYB10* homoeologs from the *F. chiloensis* ssp. *chiloensis* accessions at different allelic dosage (Supplemental Table 5). We speculate the presence of this premature stop codon might be triggering a genetic compensation response (Ma et al., 2019; El-Brolosy et al., 2019) through induction of *MYB10-1* expression (Figure 6C). Remarkably, fruits from three of the *F. chiloensis* ssp. *chiloensis* accessions (FC156, FC160 and FC187) were homozygous for the *fcmyb10-1* allele in Fvb1-2 and are completely white, not accumulating anthocyanins in the flesh or skin. In contrast, the fourth *F. chiloensis* ssp. *chiloensis* accession, FC157, was heterozygous for c. 478 G>T SNP in all *MYB10* homoeologs, and presents a light pink epidermis, indicating once more that the MYB10 C-terminal end is required to induce anthocyanin biosynthesis (Figure 6D). Notably, no color was developed in FC157 fruit flesh suggesting *MYB10* might not be expressed in this tissue.

RNA-seq data was used to analyze the expression level of the main structural genes of the anthocyanin biosynthesis pathway, finding transcript levels of most anthocyanin biosynthesis pathway genes were significantly down-regulated in all white-fleshed accessions (Figure 7). Transcripts from genes which products are involved in the early steps of the pathway, including *PAL, C4H* and *4CL* from the common phenylpropanoid pathway, and the EBGs *CHS*, *CHI* and *F3H*, were the most affected. However, the expression level of one *UFGT,* a LBG, and *GST*, were also notably reduced in the accessions with the *fcmyb10-1* allele compared to the red-fleshed ‘Senga Sengana’. *DFR* and *ANS* did not seem to be under the transcriptional control of MYB10. Even though their expression is downregulated in some of the white-fleshed or completely white accessions, it could be interpreted as a background effect. The most dramatic effect in terms of reduction of expression level was found in the accessions carrying the putative nonfunctional *fcmyb10-1* allele in homozygosis in chromosome Fvb1-2: FC156, FC160 and FC187. In these accessions, the expression of *CHS*, *F3H, UFGT* and *GST* was practically abolished, in agreement with the total white phenotype of their fruits, and thus confirming *fcmyb10-1* is a loss-of-function allele. In USA1 and FC285, downregulation of some of the structural genes of the anthocyanin biosynthesis pathway was more moderate, but it should be noted that fruits from these accessions have red epidermis and therefore accumulate anthocyanins. Similar to our results, down-regulation of *FaMYB10* did not affect *ANS* expression in octoploid strawberry (Medina-Puche et al., 2014) while lower *ANS* expression in white fruits than in red fruits was observed in *F. vesca* (Figure 2; Lin-Wang et al., 2014; Härtl et al., 2017). Therefore, FaMYB10 and FvMYB10 may differ in the regulation of *ANS* as has been previously suggested (Lin-Wang et al., 2014).

**Figure 7.**
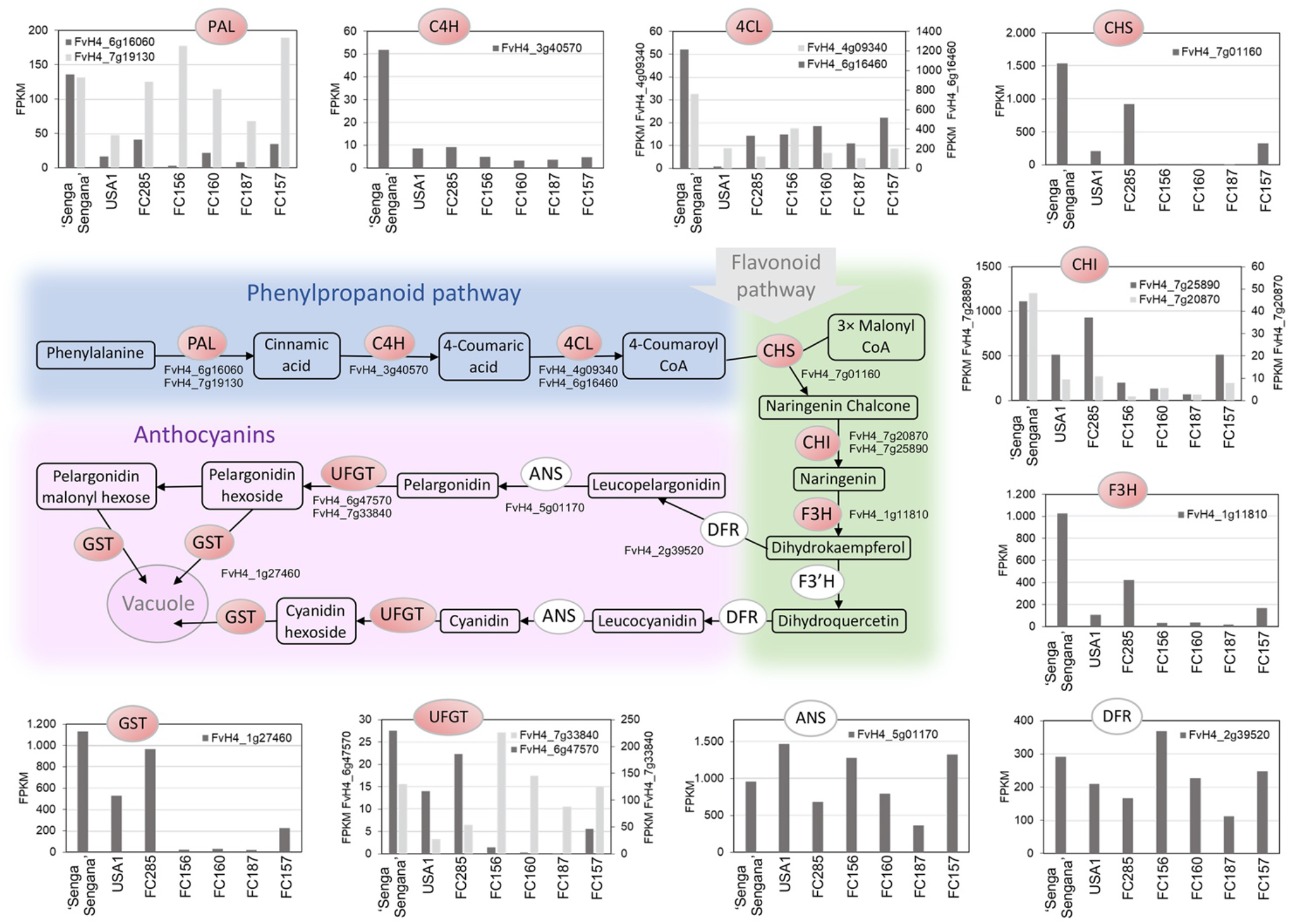
RNA-seq expression data of structural genes from anthocyanin biosynthetic pathway in ‘Senga Sengana’ and white-fleshed *F. chiloensis* accessions. FPKM values for each homoeolog of the same gene were added up and the resulting value represented only if above 10 FPKM. To avoid naming all ‘Camarosa’ homoeologs, each gene was labelled using *F. vesca* codes. Highlighted in red are those enzymes which are encoded by genes dramatically downregulated in all white fleshed accessions. FPKM values and ‘Camarosa’ gene code for each homoeolog are provided in Supplemental Table 6.

### Mining putative regulatory elements in *FaEnSpm-2*

Next, we investigated whether the *FaEnSpm-2* element could be responsible for *MYB10-2* activation in red-fleshed fruits by providing novel cis-regulatory sites behaving as enhancers, conferring responses to different stimuli and/or providing flesh-specific expression. PlantPAN 3.0 database (Chow et al., 2018) was used to interrogate the presence of putative transcription factor binding sites (TFBSs) in *MYB10-2* upstream regulatory regions from ‘Camarosa’ and the white-fleshed accessions USA2 and FC157. The ‘Camarosa’ *FaMYB10-2_pro_* sequence analyzed spanned 3,986 bp upstream of the ATG start codon, including the proximal 3 kb of *FaEnSpm-2* element and 986 bp of promoter sequence downstream of *FaEnSpm-2* insertion site. For USA2 and FC157 *MYB10-2_pro_* sequences, 2,069 bp and 2,729 bp upstream of the initial ATG were surveyed. All three fragments shared almost 1 kb of the most proximal promoter region downstream of *FaEnSpm-2* insertion point. In the overlapping region 36 SNPs and 4 small INDELs were found, not considering the large 660 bp insertion in FC157 *MYB10-2_pro_*. Results were filtered to a list of 84 putative *cis*-regulatory elements (at 156 positions) found exclusively at ‘Camarosa’ *FaMYB10-2_pro_* (Supplemental Table 7.1). We next focused our analysis on those motifs potentially relevant in the context of fruit ripening and anthocyanin biosynthesis. Among them, hormone-responsive elements such as ABA- and methyl jasmonate (MeJA)-responsive elements were significantly enriched, with a total of 13 and 4 different putative motifs, respectively (Figure 8A, Supplemental Table 7.2). Additionally, three different MYB binding motifs were identified. These elements might be of special significance as they could provide a feed-forward mechanism resulting in *MYB10* upregulation. In particular, the BOXLCOREDCPAL (ACCWWCCT) motif, a variant of the conserved MYBPLANT (MACCWAMC) and MYBPZM (CCWACC) elements, is likely to be bound by MYB10 as do other R2R3-MYBs involved in phenylpropanoid and anthocyanin biosynthesis (Sablowski et al., 1994; Grotewold et al., 1994; Jian et al., 2019). The presence of two transcription enhancers and two sugar-response elements at *FaEnSpm-2* might also be influencing *MYB10-2* expression in fruit flesh (Figure 8A, Supplemental Table 7.2). Notably, the majority of red-flesh associated TFBSs identified were located within the *FaEnSpm-2* TE, while only 6 elements (at 6 positions) specific to ‘Camarosa’ *FaMYB10-2_pro_* were detected in the ~ 1 kb promoter fragment shared by the three accessions (Supplemental Table 7.1).

**Figure 8.**
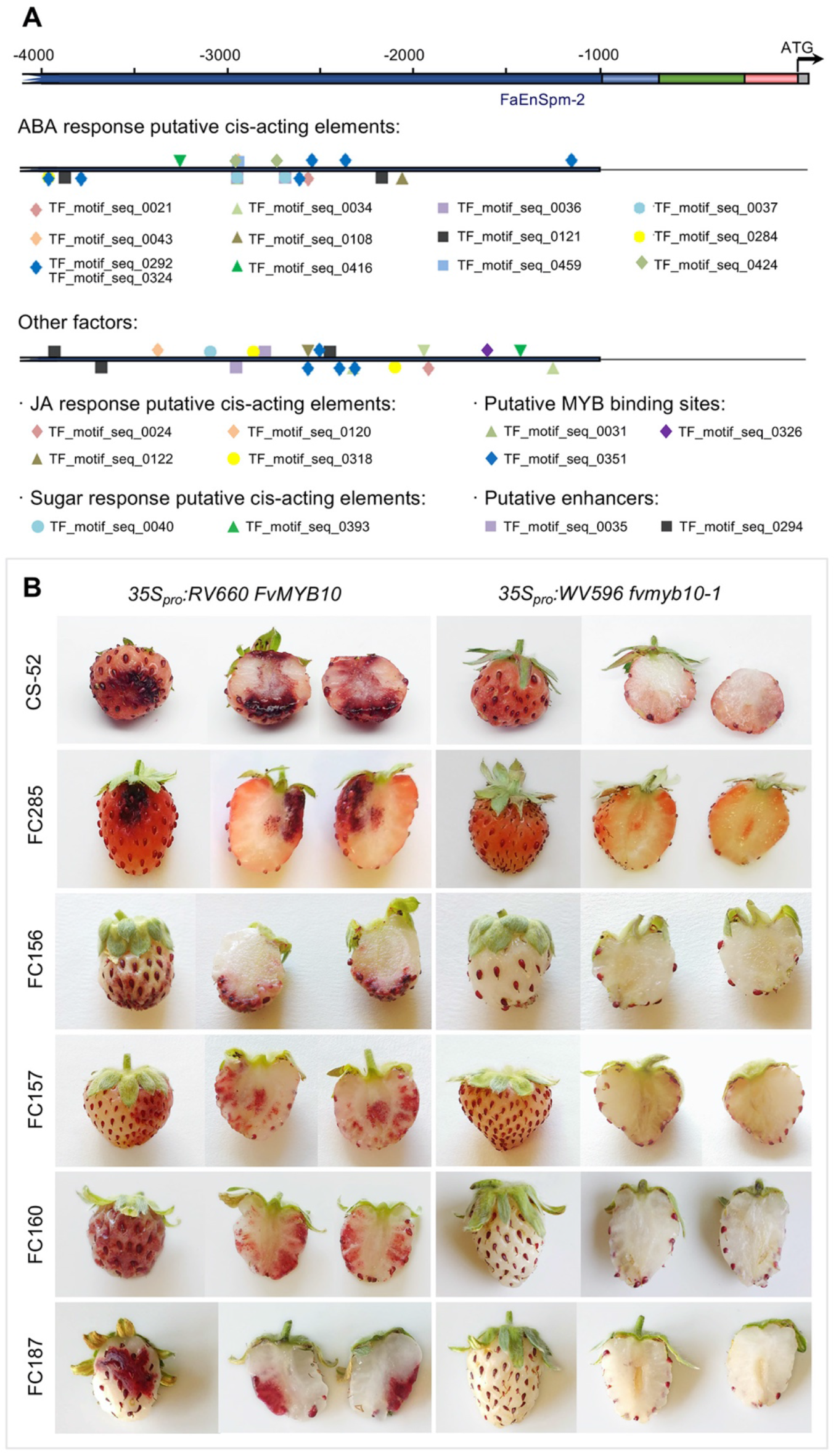
Prediction of putative regulatory cis-elements in the promoter region of ‘Camarosa’ *FaMYB10-2* not found in *MYB10-2_pro_* from the white fleshed accessions USA2 or FC157. Functional validation of *MYB10* as the causal agent under the internal fruit color QTL. **(A)** Schematic representation of selected regulatory cis-elements exclusively predicted in ‘Camarosa’ *MYB10-2_pro_* allele but not present in the white-fleshed accessions USA2 or FC157. Additional information on these elements can be found in Supplemental Table 7. **(B)** Complementation of white flesh phenotype in *F. chiloensis* accessions by transient *FvMYB10* overexpression. A representative fruit from each assay is shown.

Cultivated strawberry fruits accumulate anthocyanins in both the flesh and the skin, while it is more common to find fruits and vegetables accumulating anthocyanins only in their skin (Chaves-Silva et al., 2018; Jaakola, 2013). A wide variation in skin and flesh color has also been reported in apple and shown to be associated with different polymorphisms, including the presence of TEs, in different alleles of the apple *MYB10* ortholog (Espley et al., 2009; Chagné et al., 2013; Zhang et al., 2019). As in apple, in other species such as citrus, TEs have been shown to enhance fruit specific MYB expression (Butelli et al., 2012). In contrast, the R1 and R6 motifs found in apple *MYB10* promoters and shown to enhance *MYB10* expression in different species (Espley et al., 2009; Brendolise et al., 2017) were not detected in any of the analyzed strawberry promoter sequences.

Several studies have shown that light intensity and quality are important enhancers of anthocyanin biosynthesis, especially in fruit skin (as reviewed in Jaakola, 2013). Furthermore, *MYB10* has been shown to be a positive regulator of light-controlled anthocyanin biosynthesis in apple and strawberry (Lin-Wang et al., 2010; Li et al., 2012; Kadomura-Ishikawa et al., 2014). Independently from light, ABA also promotes *FaMYB10* expression, resulting in induction of anthocyanin biosynthesis (Medina-Puche et al., 2014; Kadomura-Ishikawa et al., 2014). Strawberry is a non-climacteric fruit and ABA is known to play a crucial role in the ripening process (Jia et al., 2011; Chai et al., 2011). Additionally, sucrose can induce ABA accumulation and promote strawberry fruit ripening by ABA-dependent and -independent mechanisms (Jia et al., 2016; 2013). In a recent study, Luo et al. (2019) showed that both ABA and sugar play a synergistic role in promoting strawberry fruit ripening, including anthocyanin accumulation. Finally, methyl jasmonate (MeJA) treatment has also been shown to increase anthocyanin accumulation in strawberry fruit (Concha et al., 2013). Further work will be required in order to better understand the significance of the predicted putative cis-elements and the molecular mechanisms driving higher *MYB10* expression level in red-fleshed accessions. However, it is tempting to speculate that in the interior of the receptacle, where light quality or intensity might not be as effective in inducing *MYB10* expression, other endogenous signals, such as ABA, sucrose or MeJA, would be required in order to accumulate enough *MYB10* transcript level to induce anthocyanins light-independently. In this scenario, promoters lacking the *FaEnSpm-2* element would not contain the putative regulatory sequences able to recruit the transcription factors responding to those stimuli and would fail to induce anthocyanin biosynthesis in the receptacle flesh.

Known *F. vesca* accessions are characterized by red skin and white flesh, while cultivated strawberry cultivars display a characteristic and preferred red interior. Like in the white-fleshed *F. chiloensis* accessions analyzed in this study, *F. vesca MYB10_pro_* lacks a *FaEnSpm-2-like* TE, as it is not predicted in the *F. vesca* Hawaii4 reference genome (Shulaev et al., 2011) and was not detected by PCR in accessions analyzed in this study as RV660 or WV596 (data not shown). When putative TFBS were predicted in the homologous *MYB10_pro_* region from *F. vesca,* 941 bp long, only 5 of the 84 elements exclusively found at ‘Camarosa’ *FaMYB10-2_pro_* (Supplemental Table 7.3) were detected. None of them were among the selected cis-elements (Supplemental Table 7.2) potentially relevant for fruit anthocyanin production.

### Transient overexpression of *MYB10* overcomes white flesh and skin phenotypes in strawberry fruit

Finally, if reduced *MYB10* expression in the interior of the receptacle is the underlying cause of white-fleshed phenotype, it is expected that increasing *MYB10* dose should lead to phenotype complementation. Thus, functional validation was performed in fruits from CS-52, a white-fleshed F_2_ line from SS×FcL population, and the rest of white-fleshed *F. chiloensis* accessions FC156, FC157, FC160, FC187 and FC285. In all of them, transient expression of *FvMYB10* under the control of the CaMV 35S promoter was able to promote anthocyanin accumulation in both fruit flesh and epidermis (Figure 8B).

Taken together, our results confirm *MYB10-2*, the *MYB10* homoeolog from the *F. iinumae*-derived subgenome, as the dominant homoeolog in octoploid strawberry and, furthermore, the causal locus responsible for natural variation in internal and external fruit color. Alleles from this locus bearing the *FaEnSpm-2* CACTA element in the upstream regulatory region are associated with enhanced *MYB10* expression, which results in anthocyanin accumulation in the inner receptacle. We postulate that the increase in *FaMYB10* expression might be due to an expansion of its expression domain into fruit flesh. Additional analysis of *FaEnSpm-2* indicated the presence of putative promoter motifs involved in ABA, MeJA and sugar response, as well as predicted MYB binding sites potentially involved in a positive feedback mechanism. Along with the promoter polymorphism, a number of *F. chiloensis* ssp. *chiloensis* accessions with white flesh and skin carry the novel *fcmyb10-1* allele at all three full length homoelogous copies of *FcMYB10,* although at different doses (Supplemental Table 5). The predicted fcmyb10-1 is a truncated protein lacking 74 residues from the end portion of the activation domain. We have shown fcmyb10-1 fails in activating downstream anthocyanin structural genes *PAL, C4H, CHS, F3H and GT* and leads to completely white or pink fruits depending on allelic dosage (Figure 7). An independent missense mutation, *famyb10-1,* was also found in this study for lines with white skin color from UF breeding population 17.66. The mutation is same as a previously identified INDEL (Wang et al., 2020). Our study precisely located this mutation to the dominant homoeologous allele on Fvb1-2 and showed that it controls fruit skin color. Whereas polymorphisms found in the coding region seem to be more specific to a subset of accessions, the promoter polymorphism described in this study has been shown to be common to a taxonomically diverse selection of white-fleshed accessions.

### Development of predictive high-throughput markers for fruit flesh and skin color

Development of high-throughput DNA tests with direct applicability for strawberry breeders has lagged behind other crops due to the complexity of the octoploid strawberry genome and the lack of quality subgenome-scale sequence information. Nevertheless, progress in that direction is expected since the recent release of the first high-quality chromosome-scale octoploid strawberry genome (Edger et al., 2019a). Assays for SNP detection such as kompetitive allele-specific polymerase chain reaction (KASP; Semagn et al., 2013) and high-resolution melting (Wittwer et al., 2003) have become tests of choice for breeding applications due to accuracy, ease of scoring, and applications to polyploid species such as strawberry (Whitaker et al., 2020).

We first developed an HRM marker (WS_CID_01) to predict the presence of the *famyb10-1* allele for the 8-bp INDEL in *MYB10* using the 17.66 population (102 individuals). Marker WS_CID_01 perfectly predicted white and red skin color (Figure 9A, Supplemental Table 8). For flesh color prediction the IFC-1 marker we developed accurately predicts *MYB10-2* alleles and flesh color phenotypes. However, to develop a more high-throughput assay for marker-assisted selection of white- or red-fleshed strawberries, we designed the KASP marker IFC-2. To identify homoeolog-specific primers that were not expected to amplify *MYB10* homeologs in other sub-genomes or other off-target DNA sequences, the promoter fragments here described were aligned to those of ‘Camarosa’ and other octoploid accessions from Hardigan et al. (2020). We identified an A/G SNP 20 bp upstream of the initial ATG and 966 bp downstream of the *FaEnSpm-2* insertion (Supplemental Figure 9). The A allele for the IFC-2 marker was exclusively observed in white-fleshed individuals and accessions. The IFC-2 KASP marker was used to genotype the SS×FcL mapping population (*n* = 108), in addition to two red-fleshed *F. ×ananassa* cultivars (‘Camarosa’ and ‘Candonga’), the red-fleshed *F. chiloensis* accession FC154, and the 6 white-fleshed *F. chiloensis* accessions described earlier. The IFC-2 KASP marker produced codominant genotypic clusters and was 99% predictive of white- and red-fleshed phenotypes (Figure 9B). The red-fleshed allele (G) was observed in the white-fleshed accession FC285, which was the only disparity in our study. The region targeted with this marker is highly polymorphic and even though two common primers accounting for an additional SNP were employed, it might not work in some specific backgrounds. Still, the fact it worked in > 99% of the genotypes tested, makes this marker a valuable diagnostic tool for breeders. Both, the WS_CID_01 and IFC-2 markers should facilitate the rapid introgression of target alleles into elite backgrounds and accelerate the development of white or red-fleshed cultivars, respectively. rate.

**Figure 9.**
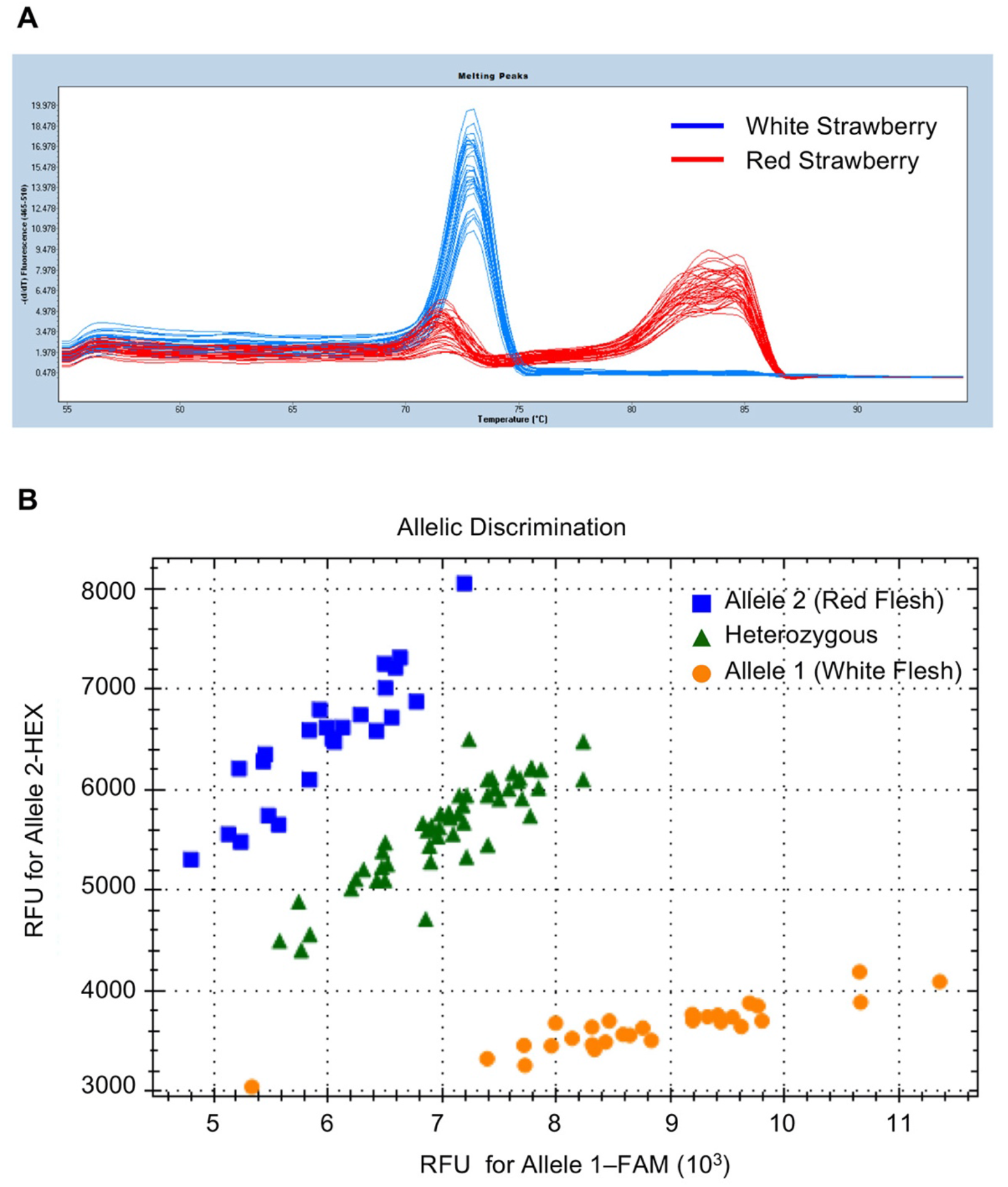
High-throughput markers for fruit skin and flesh color. **(A)** HRM analysis of WS_CID_01 marker associated with white fruit color using 102 breeding accessions. The blue and red peaks are associated with white and red color in strawberry, respectively. **(B)** Scatter plots of IFC-2 KASP assay in octoploid strawberry accessions reveal the clustering of white-fleshed lines on X-(FAM) axis (yellow dots), homozygous red-fleshed lines on Y-(HEX) axis (blue squares) and heterozygous lines in between (green triangles).

## CONCLUSIONS

In this study we have analyzed a broad range of strawberry accessions belonging to the diploid *F. vesca* species as well as the octoploids *F. chiloensis* and *F*. ×*ananassa*. Cultivated strawberry, *F. ×ananassa* is an allopolyploid derived from an interspecific cross between *F. chiloensis* and *F. virginiana.* Fruits from *F. vesca* are characterized by having white interior and red skin, however several natural mutants can be found which do not accumulate anthocyanins in fruits and have white/yellowish skin as well. We took advantage of this natural variation to explore the possibility of finding other factors besides MYB10 involved in fruit pigmentation. A total of 12 white skin accessions from different geographical origins were analyzed and in all 12 the lack of anthocyanins was explained solely by multiple independent mutations in the *FvMYB10* ORF. We were able to identify two new alleles *(fvmyb10-2* and *fvmyb10-3)* and a large chromosomal deletion in Fvb1 removing the *FvMYB10* gene.

Polyploidy, or whole genome duplication, is an important contributor to speciation in flowering plants. The formation of an allopolyploid involves the merger of genomes with separate evolutionary histories and often brings along different mechanisms to compensate for the increased gene dosage, including subgenome dominance. One important factor for subgenome dominance is a lower methylated TE abundance relative to other subgenomes (Alger and Edger, 2020). Strawberry is a complex allopolyploid that exhibit dominance in the *F. vesca-*derived subgenome, which contains the lowest TE density (Edger et al., 2019a). In particular, for anthocyanin biosynthesis, chromosomes derived from the *F. vesca* subgenome are responsible for expression of 88% of structural genes of the pathway (Edger et al., 2019a). However, in this study we demonstrated that anthocyanin biosynthesis is activated predominantly by *MYB10-2,* the homoeolog lying in the *F. iinumae-derived* subgenome. We have shown the expression dominance of *FaMYB10-2* in two strawberry cultivars, ‘Camarosa’ and ‘Senga Sengana’, both with deep red flesh. High expression of *MYB10-2* in fruits and red flesh color were strongly associated with the presence of the CACTA element *FaEnSpm-2* about 1 kb upstream of the ORF in all octoploid accessions included in this study. In contrast, strawberry accessions lacking the TE were characterized by a white flesh, while heterozygous lines displayed an intermediate phenotype indicating incomplete dominance, as also shown in the QTL analysis where the dominance effect of *qFleshCol-1-2* QTL was estimated at 0.24-0.36, depending on the year (Supplemental Table 4).

Since cultivated strawberry breeding began in the 1800s using a small number of lines (Darrow, 1966), the diversity of *F. ×ananassa* has been increased and its genome reshaped by repeated interspecific hybridization with phylogenetically diverse *F. chiloensis* and *F. virginiana* accessions (Darrow, 1966; Gil-Ariza et al., 2009; Hardigan et al., 2018; Liston et al., 2014; Hancock, 1999; Bringhurst and Voth, 1984; Hancock et al., 2002; 2001). Introgression of beneficial alleles from these species also results in the accumulation of unfavorable alleles in cultivated strawberry. One unwanted trait in most strawberry breeding programs developing varieties with red skin is white internal flesh. On the other hand, strawberries with both white skin and white flesh are becoming popular in some countries such as Japan. Furthermore, current breeding programs are particularly directed to increase fruit quality and pathogen resistance (Whitaker et al., 2020), and *F. chiloensis,* as *F. virginiana,* represent a reservoir of interesting alleles for future improvement of biotic and abiotic stress tolerance and fruit flavor and aroma (Aharoni et al., 2004; Johnson et al., 2014; Hancock, 1999). Our findings showed a simple genetic control of *MYB10* over strawberry skin and flesh color, making it a good target for molecular breeding. Therefore, we anticipate that markers developed in this study will enable efficient breeding advances, particularly when wild relatives are used as parental lines.

## METHODS

### Plant Materials and Phenotypic Evaluation

#### Diploid *Fragaria vesca* Germplasm

An F_2_ mapping population of 145 lines between the everbearing, non-runnering *F. vesca* ‘Reine des Vallées’ (ESP138.660; RV660) and the white-fruited *F. vesca* ESP138.596 (WV596) was developed from one F_1_ plant (F_1_-014) that was able to produce runners and had red fruits. The population was grown in greenhouse conditions in Málaga (Spain) and phenotyped for two years, and the two traits, runnering and the red/white fruits, were found to segregate independently as two single mutations. The rest of *F. vesca* accessions studied were grown in the same conditions and are described in Supplemental Table 3 and include ‘South Queen Ferry’, GER1 and GER2 from the “Professor Günter Staudt collection” (Dresden, Germany), and UK13, SE100 and FIN12 from Dr. T. Hytönen and Dr. D. Posé collection.

### Octoploid *Fragaria* Germplasm

The University of Florida breeding population 17.66 was derived from a cross between FL 13.65-160 (red) and FL 14.29-1 (white) selections (Supplemental Figure 4; Supplemental Table 3 and 8). The population (102 individuals) was grown in open field conditions with two plants per plot, and fruit skin color was assayed three times from December 2018 to February 2019 at the UF/IFAS Gulf Coast Research and Education Center (Balm, Florida).

To generate the SS×FcL F_2_ population of 105 progenies, a cross between *Fragaria ×ananassa* cv. ‘Senga Sengana’ and *F. chiloensis* ssp. *lucida* USA2 was performed at Hansabred, Germany in 2008 (Supplemental Figure 5). An F_1_ seedling, cloned under the number P-90999, was selected among the progeny based on key breeding traits such as tolerance to two spotted spider mite *(Tetranychus urticae* Koch), yield, and fruit aroma and color. The selected F_1_ individual was self-pollinated to obtain the F_2_ mapping population. The population was grown under field conditions at Hansabred and scored for fruit skin, flesh and core color in three seasons (2014, 2016 and 2019). Skin color was evaluated using a five-score scale from completely white to dark red. Flesh and core color was evaluated using a three-score scale: 1, white; 2, light red; and 3, red (Supplemental Figure 10). The parental line *F. chiloensis* ssp. *lucida* USA2 is a male individual that does not set fruit, and therefore, USA1, a ‘sister’ female plant collected from the same area was included in phenotypic and molecular analyses, as other *F. chiloensis* accessions from diverse origins (Supplemental Table 3).

### DNA Extraction and QTL-Seq Analysis in Diploid *Fragaria vesca*

DNA was extracted from young leaf of parental, F_1_ and each F_2_ lines using the CTAB method (Doyle and Doyle, 1990) with minor modifications. For rapid mapping of the fruit color mutation, we performed whole-genome resequencing of DNA from two bulked populations as previously described (Takagi et al., 2013). The two pools, red fruit pool (RF) and white fruit pool (WF), were produced by mixing an equal amount of DNA from 34 RF and 32 WF F_2_ lines, respectively. Pair-end sequencing libraries with insert sizes of approximately 350 bp were prepared and 100 bp length sequences produced with a 50x genome coverage using an Illumina HiSeq 2000. The reads from RF and WF pools were mapped independently against the last version of the *Fragaria vesca* reference genome, F_vesca_H4_V4.1 (Edger et al., 2018). The low-quality reads (< Q20 Phred scale) were filtered using samtools method (Li et al., 2009). Then, duplicates originated from polymerase chain reaction were eliminated using picard-tools program. For single nucleotide variants (SNV) and small INDELs calling process, the gatk algorithm (McKenna et al., 2010) was applied. The large INDELs were identified using Manta algorithm (Chen et al., 2016). The variants with coverage less than 20 reads in both samples were not considered in down-stream analyses.

SNP frequencies in each pool were calculated as the proportion of reads harboring the SNP different from the reference genome. Thus, SNP frequency = 0 if all short reads match the reference sequence and 1 if they correspond to the alternative allele. SNP positions with read depth <7 and frequencies <0.3 in both pools were excluded, as they may represent spurious SNPs called due to sequencing or alignment errors. The parameter ΔSNP-index was calculated as the absolute difference between SNP frequencies in the RF and WF pool samples. For detection of significant genomic regions, the genome was split in 3-Mb sliding windows with of 10 kb increments. For each window, the average ΔSNP-index was calculated and used for the sliding window plot. For identification of statistically significant regions, a Wilcoxon test was applied using a confidence *P*-value of 0.03.

### Whole Genome Resequencing of FIN12 and Data Analysis

Whole genome resequencing of FIN12 accession was carried out at DNA Sequencing and Genomics Laboratory, Institute of Biotechnology, University of Helsinki, Finland using Illumina NextSeq 500 sequencer. Library preparation, pairend sequencing (150 + 150 bp), and the alignment of the sequencing reads on the F. vesca H4 reference genome v4.1 (Edger et al., 2018) was carried out as described earlier (Koskela et al., 2017). Sequencing data is stored at NCBI Short Read Archive (https://www.ncbi.nlm.nih.gov/sra) under the accession number XXXXXX.

### GWAS Analysis in *F. ×ananassa* Breeding Germplasm

A total of 95 individuals from UF family 17.66 were used for a genome-wide association study. Total genomic DNA was isolated from leaf tissues using a modified CTAB method described by (Noh et al., 2018). A whole-genome SNP Genotyping was performed using the FanaSNP 50K Array (Hardigan et al., 2020). A general linear model (GLM), mixed linear model (MLM), and multi-locus mixed-model (MLMM) were used for association tests using GAPIT v2 performed in R (Tang et al., 2016; Team, 2014; Liu et al., 2016). Manhattan plots were created using the R package qqman version 0.1.4 (Turner, 2014).

### Linkage Mapping and Detection of QTL in Octoploid *Fragaria*

DNA was extracted from young leaf of parental, F_1_ and the 105 F_2_ lines using the same CTAB method as previously described for diploid *Fragaria.* A total of 16,070 SNP markers were produced using the strawberry DArTseq platform (Sánchez-Sevilla et al., 2015). SNP markers that were monomorphic in the progeny were removed, as markers that did not fit the expected 3:1 segregation using the χ^2^ test (p =0.05). Next, markers with rowsums < 400 were also removed. Finally, markers with more than 5% missing scores (more than five progeny lines) were excluded, resulting in a total of 9,005 dominantly scored SNP markers. For 5,856 markers, the reference and the alternative allele segregated in a 1:2:1 ratio (as expected for a F_2_ population) and were transformed into 2,523 codominant markers (COD-SNPs). The 2,523 COD-SNPs were used together with 2,446 dominant SNPs (DOM-SNPs) for mapping using JoinMap 4.1 (van Ooijen, 2006). First, JoinMap software was used coding the markers as CP population to infer the phase of those markers heterozygous in both parental lines and thus with unknown origin in the F_1_ line. Phase information was then used to assign phases and to code the SNP markers as a F_2_ population. Grouping was performed using independence LOD and the default settings in JoinMap 4.1 and linkage groups were chosen at a LOD of 9 for all the 28 groups obtained. Map construction was performed using the maximum likelihood mapping algorithm and the following parameters: Chain length 5,000, initial acceptance probability 0,250, cooling control parameter 0,001, stop after 30,000 chains without improvement, length of burn-in chain 10,000, number of Monte Carlo EM cycles 4, chain length per Monte Carlo EM cycle 2,000 and sampling period for recombination frequency matrix samples: 5. A total of 422 identical loci and 517 loci with similarity >0.99 were removed. The seven HGs were named 1 to 7, as the corresponding LGs in the diploid *Fragaria* reference map. SNP marker sequences were blasted to the recently published ‘Camarosa’ genome and assigned to the best matching chromosome. LGs within homoeologous groups were then named 1 to 4 according to the corresponding ‘Camarosa’ chromosome number (Edger et al., 2019a).

QTL analyses were performed using MapQTL 6 (van Ooijen, 2009). The raw relative data was analyzed first by the nonparametric Kruskal-Wallis rank-sum test. A stringent significance level of *p* ≤ 0.005 was used as a threshold to identify markers linked to QTL. Second, transformed data sets for non-normally distributed traits were used to identify and locate mQTL using Interval Mapping (IM) with a step size of 1 cM and a maximum of five neighboring markers. Significance LOD thresholds were estimated with a 1,000-permutation test for each trait. The most significant markers were then used as co-factors for restricted multiple QTL mapping (rMQM) analysis. mQTL with LOD scores greater than the genome-wide threshold at *p* ≤ 0.05 were declared significant. Linkage maps and QTLs were drawn using MapChart 2.2 (Voorrips, 2002).

### RNA Extraction and RT-qPCR

Total RNA was extracted from three independent replicates of two pools of ripe fruits (WF and RF) from the same F_2_ lines of ‘RV660’ x ‘WV596’ population used for QTL-seq. A modified CTAB method (Gambino et al., 2008) with minor modifications was followed. Basically, 300 mg of powdered fruit sample was used as starting material and RNA was resuspended in 30 μL sterile water. After digestion with TURBO DNA-free™ Kit (Thermo Fisher Scientific AM1907), cDNA was synthesized from 1 ug of RNA using the High-Capacity cDNA Reverse Transcription Kit (Thermo Fisher Scientific 4368814). RT-qPCR analysis was performed with iQ SYBR Green Supermix in an iCycler iQ5 system (Bio-Rad) following the manufacturer’s instructions. Three technical replicates were performed for each biological replicate. Relative expression of each gene was calculated using GADPH as internal reference gene. Sequences of primers used in this study are listed in the Supplemental Table 9.

### Semi-polar Metabolite Analysis Using UPLC-Orbitrap-MS/MS

To analyze the effect of *FvMYB10* mutation, two bulked pools of ripe fruits from the same F_2_ lines used for QTL-seq and RT-qPCR were prepared in triplicate. Fruit tissue from each pool was grinded in liquid nitrogen and stored at −80°C. Secondary metabolite profiles of RF and WF pools were performed using methods described by (Vallarino et al., 2018) using a Waters Acquity UPLC system. Full documentation of metabolite profiling data acquisition and interpretation is provided in Supplemental Table 2.

### Quantification of Fruit Quality Parameters

For the determination of SSC, TA, TEAC and ascorbic acid, frozen fruit powder from the same samples (in biological triplicate) used for expression and metabolomic studies were used. For SSC, approximately 0.5 g of sample was defrosted to room temperature and the puree deposited onto the lens of a refractometer (Atago PR32, Japan). Titratable acidity was evaluated with an automatic titrator (Titroline easy, Schott North America, Inc.) as previously described (Zorrilla-Fontanesi et al., 2011). Polyphenols were extracted in 80% methanol and antioxidant capacity of fruit samples was measured by the ability of antioxidant molecules to quench the ABTS^·+^ radical cation [2,2’-azinobis(3-eth-ylbenzothiazoline-6-sulfonate); Sigma-Aldrich] in comparison with the antioxidant activity of standard amounts of Trolox. The Trolox equivalent antioxidant capacity (TEAC) assays were performed as described (Pellegrini et al., 1999; Re et al., 1999) using 2 μL of sample and 250 μL of radical reagent. The absorbance at 734 nm was measured after 5 min at 25°C in a spectrophotometer. Results were expressed in μmoles of Trolox equivalents per gram of fresh weight (μmol TE/g FW).

L-Ascorbic acid (L-AA) was measured by HPLC (Davey et al., 2006; Zorrilla-Fontanesi et al., 2011). In brief, 0.25 g of fruit powder was homogenized with 1 ml cold extraction buffer (metaphosphoric acid 2%, EDTA 2 mM) and kept at 0°C for 20 min. Then, samples were centrifuged at 14,000 rpm for 20 min at 4°C and the supernatant was filtered (0.45 mm membrane) and transferred to HPLC vials kept on ice. Finally, 5 ml was injected in a Rx-C18 reverse-phase HPLC column (4.6 x 100 mm, 3.5 mM, Agilent Technologies) and detection was carried out at 254 nm in a photodiode array detector (G1315D, Agilent Technologies). The mobile phase consisted of a filtered and degassed solution of 0.1 M NaH2PO4 0.2 mM EDTA pH 3.1 (Harapanhalli et al., 1993) and a flow rate of 0.7 ml/ min. L-AA content was calculated by comparison with values obtained from a standard curve and were expressed as mg L-AA per 100 g of fresh weight.

### Amplicon Sequence Analysis

Amplicon sequencing was performed using cDNA of white and red fruit accessions from family 17.66. The fruits from each white-skinned and red-skinned accessions were collected in the field, and two millimeters thickness of fruit skin was collected using surgical blade for RNA extraction. Total RNAs were extracted with RNeasy^®^ Mini Kit (Qiagen, Germany) and cDNA synthesis was carried out using LunaScript™ RT SuperMix Kit (New England Biolabs, USA), following the manufacturer’s instructions. For amplicon sequencing, primer sets were developed for the coding region of *MYB10* gene (cv. Camarosa, maker-Fvb1-2-snap-gene-157.15-mRNA-1) using IDT’s PrimerQuest Software (Supplemental Table 9). PCR products were checked in 3% agarose gel, and further purified for amplicon sequencing at Genewiz (South Plainfield, NJ). The generated sequencing reads from each white and red fruit pool (10 samples per each pool) were mapped to the *MYB10* gene, maker-Fvb1-2-snap-gene-157.15-mRNA-1, using default values in Geneious v.11.0.5. The consensus sequence of *MYB10* coding region of white and red strawberry pools were aligned using T-coffee (Notredame et al., 2000).

### Gene and Promoter Cloning and Sequence Homology Analysis

Gene and promoter amplification were performed using MyFi™ DNA Polymerase (Bioline). FvMYB10-*gypsy* and Fvb1 FIN12 genomic fragments flanking the deleted region were both amplified with 5Prime PCR Extender System (5Prime) using the provided 10X Tunning Buffer and following the manufacturer instructions.

PCR products <8kb were purified with FavorPrep GEL/ PCR Purification Kit (Favorgen). If >8kb, PCR products were precipitated (30% PEG8000; 30mN MgCl2). Purified products were cloned into the pGEM^®^-T Easy Vector Systems (Promega) for Sanger sequencing.

Multiple sequence alignment of *MYB10* promoter fragments was performed using Geneious algorithm (Gap open penalty=30; Gap extension penalty=0) and used for homology tree construction (Genetic distance model: Jukes-Cantor; Tree build method: Neighbor-Joining), both in the Geneious platform (Kearse et al., 2012).

### RNA-Seq Analysis, Differential Gene Expression and Variant Calling

Total RNA was extracted from biological replicates of ripe fruits from the seven selected octoploid accessions as described above. RNA quantity, quality and integrity were determined based on absorbance ratios at 260 nm/280 and 260 nm/230 nm using a NanoDrop spectrophotometer (ND-1000 V3.5, NanoDrop Technologies, Inc.), by agarose gel electrophoresis, and further verified using a 2100 Bioanalyzer (Agilent, Folsom, CA). Sample RIN values ranged between 7.4 and 8.4. Libraries were produced following Illumina’s recommendations at Sistemas Genómicos facilities, where they were sequenced by pair-end sequencing (100 bp × 2) in an Illumina Hiseq 2500 sequencer. Over 200.4 million reads were generated and filtered for high-quality reads by removing reads containing adapters, reads shorter than 50 bp and reads with Q-value ≤ 30 using Fastq-mcf from ea-utils (Aronesty, 2011) with parameters -q 30 -l 50. Following analyses were based on processed clean reads.

Read mapping, differential gene expression analysis and clustering of seven selected octoploid accessions and previous ‘Camarosa’ tissues experience (Sánchez-Sevilla et al., 2017) were made with “Tuxedo suit” (Hisat2/Cufflinks/CummRbund) according to standard procedures (Trapnell et al., 2012). As reference genome (Edger et al., 2019a) we used the latest assembled *F. ×ananassa* genome (‘Camarosa’ Genome Assembly v1.0.a1: F_ana_Camarosa_6-28-17.fasta) and annotation (Fxa_v1.2_makerStandard_MakerGenes_woTposases.gff). Files were downloaded from the GDR web site (https://www.rosaceae.org/). Overall alignment rate was 90.72%.

Variant calling was performed using the haplotype-based variant detector FreeBayes v1.0.2-16-gd466dde-dirty (Garrison and Marth, 2012). To identify the different variants at each position in the reference *F. ×ananassa* genome, alignment BAM files of the different accessions were labeled with samtools. VCF file was generated with FreeBayes with all parameters set default. All bioinformatic processes were performed at the Picasso cluster facilities at SCBI, Málaga (http://scbi.uma.es).

### Transient Over-expression by Agro-infiltration of Fruit

Transient expression studies in strawberry fruits were performed as described (Hoffmann et al., 2006) with minor modifications. Full-length wildtype *FvMYB10* and the truncated *fvmyb10-2* cDNAs were amplified from total cDNA from ripe fruits of F_2_ RV660 × WV596 population RF or WF pools respectively. Primers FvMYB10-attB1 F and FvMYB10-attB2 R were used to amplify wildtype *FvMYB10* and FvMYB10-attB1 F and ccm29 for *fvmyb10-2* amplification (Supplemental Table 9). Primer ccm29 is complementary to a region of the retrotransposon 45-70 bp downstream of the insertion point, which includes the first stop codon generated after the insertion. Each gene version was cloned into the Gateway transfer vector pDONR221 and then into the binary vector pK7WG2D under the control of the CaMV 35S promoter (Gateway BP and LR Clonase II Enzyme mix, Invitrogen). The resulting plasmids were introduced into the *Agrobacterium* strain AGL0 following the protocol of (Höfgen and Willmitzer, 1988). Immature fruits in the late green/early white stage were used for agro-infiltration using 1 ml suspension of each culture in combination (1:1) with a suspension of *Agrobacterium* LBA4404 with the p19 suppressor of gene silencing (Voinnet et al., 2003). Agro-infiltrated fruits were labelled and left to ripen *in planta* for about 7-10 days.

### HRM Marker Development and Marker Data Analysis

The primer set targeting the 8-bp mutation was designed using IDT’s PrimerQuest Software and ordered from IDT (San Jose, CA, USA) (Supplemental Table 9). PCR amplification was performed in a 5 μl reaction containing 0.5 μM of each primer set, 2× AccuStart^TM^ II PCR ToughMix^®^ (Quantabio, MA, USA), 1× LC Green^®^ Plus HRM dye (BioFire, UT, USA), and 0.5 μL of DNA. PCR conditions were as follows: an initial denaturation at 95°C for 5 minutes; 45 cycles of denaturation at 95°C for 10 seconds, annealing at 62 °C for 10 seconds, and extension at 72°C for 20 seconds. After PCR amplification, the samples were heated to 95°C for 1 minute and cooled to 40°C for 1 minute. Melting curves were obtained by melting over the desired range (60-95°C) at a rate of 50 acquisitions per 1°C. The HRM assay was performed with LightCycler^®^ 480 System II (Roche Life Science, Germany). The HRM data was analyzed using the Melt Curve Genotyping and Gene Scanning Software (Roche Life Science, Germany). Analysis of HRM variants was based on differences in the shape of the melting curves and in Tm values.

### SNP Genotyping using Kompetitive Allele-Specific PCR (KASP)

A SNP in *MYB10-2* at position −20 from the initial ATG associated with white/red flesh color in the sequenced accessions was converted into a KASP assay. In order to produce a sub-genome specific assay, a combination of four primers was used in each reaction, two common forward primers (ccm90 and ccm91) to account for a background-specific G/A SNP and two allele-specific primers (ccm92 and ccm93) each carrying the standard FAM™ or HEX™ dye tails and the targeted SNP at the 3’ end. Generally a single common primer is used for standard KASP assays but the high degree of polymorphism in the targeted region made necessary to include an extra common forward primer to account for a different, background associated, G/A SNP not linked with the white-fleshed trait. Reaction volumes of 5 μL for 384-well plates were used. The 2.5 μl of mastermix and 0.07 μl of assay/primer mix and 2.5 μl of DNA (12.5 ng total) were used for the genotyping. Final primer concentrations in the reaction mix were: 210 nM for ccm 90 and ccm91; 150 nM for ccm92 and 190 nM for ccm93. PCR was conducted in a Biorad CFX-384 instrument using the following protocol: An initial denaturation step at 94 °C for 15 min, 12 touchdown cycles (94°C for 20 seconds, 65°C for 80 seconds, dropping 0.6°C per cycle), 29 cycles (94°C for 20 seconds, 58°C for 80 seconds) and a final recycling for 2 cycles (94°C for 20 seconds, 60°C for 80 seconds). Biorad software was used to estimate the final results.

### Accession Numbers

The raw reads data of RF and WF pools and RNA-seq have been stored at the Sequence Read Archive (SRA) of the European Nucleotide Archive (ENA) with the project reference PRJEB38128. *F. vesca* mutant *MYB10* sequences and *F. ×ananassa* promoter sequences can be found in the EMBL/GenBank data libraries with the following accession numbers: ESP138.596 *FvMYB10* ORF (xxx); GER1, GER2 and SE100 *FvMYB10* ORF (xxx); *F. chiloensis ssp. lucida* USA2 ORF (xxx), *F. chiloensis ssp. lucida* USA2 promoter fragment (xxx),…

## Supplemental data

**Supplemental Figure 1.** TEAC, SSC, TA, vitamin C in RF and WF pools.

**Supplemental Figure 2.** Transposon insertion co-segregates with the white phenotype in the entire RV660 x WV596 population.

**Supplemental Figure 3.** Large deletion in FIN12 affects ~ 100 kb and 7 predicted transcripts.

**Supplemental Figure 4.** Fruit skin color in the UF population 17.66 (FL 13.65-160 x FL 14.29-1).

**Supplemental Figure 5.** Fruit skin and flesh color phenotypes in the Hansabred SS×FcL population.

**Supplemental Figure 6.** ‘Senga Sengana’ x *F. chiloensis* ssp. *lucida* linkage map and comparison to the physical positions in *F. vesca* reference genome.

**Supplemental Figure 7.** Expression of ‘Camarosa’ *FaMYB10* homoeologs in different tissues and during fruit ripening.

**Supplemental Figure 8.** Sequence alignment of WT and mutant MYB10-2 in the UF breeding population 17.66.

**Supplemental Figure 9.** *MYB10* upstream region alignment showing the SNP targeted for KASP genotyping.

**Supplemental Figure 10.** Fruit color scale used to phenotype octoploid accessions. (**A**) Skin color scale (1-5). (**B**) Flesh color scale (1-3). Examples used in B had a score 4 in skin color. CS-63, CS-79 and CS-98 are different F_2_ lines from SS×FcL cross.

**Supplemental Table 1.** List of significant SNPs and INDELs in the RV660 × WV596 population.

**Supplemental Table 2.** Complete metabolite data.

**Supplemental Table 3.** List of *Fragaria* ssp. included in this study and description of detected mutation on *MYB10.*

**Supplemental Table 4.** QTLs detected for fruit color traits in the ‘Senga Sengana’ × *Fragaria chiloensis* ssp. *lucida* F_2_ population by Kruskal-Wallis (KW) and restricted multiple QTL mapping (rMQM) analysis.

**Supplemental Table 5.** Polymorphisms at *MYB10 homoeologs from F. chiloensis* accessions and ‘Senga Sengana’ *vs.* ‘Camarosa’ reference genome.

**Supplemental Table 6.** Expression of anthocyanin pathway genes in ‘Senga Sengana’ and white fleshed *F. chiloensis* accessions.

**Supplemental Table 7.** Predicted putative cis-elements specific of ‘Camarosa’ *FaMYB10-2_pro_*.

**Supplemental Table 8.** Phenotype and WS_CID_1 marker genotype results of the UF breeding population 17.66 (FL 13.65-160 × FL 14.29-1).

**Supplemental table 9.** List of primers.

## ACKNOWLEDGEMENTS

The authors thankfully acknowledge the computer resources and the technical support provided by the Plataforma Andaluza de Bioinformática of the University of Málaga. We thank Francisco J. Durán and the UF/IFAS strawberry breeding team for his excellent care of strawberry germplasm. This work was supported by grants RFP2015-00011-00-00 (Spanish Ministry of Economy and Competitivity and FEDER), PR.AVA.AVA2019.034 (IFAPA, FEDER funds), ERC-2014-StG 638134 (European Research Council), and by the European Union’s Horizon 2020 research and innovation programme (GoodBerry; grant agreement number 679303). This study is also part of the joint research network SPIRED which was funded by the German Federal Ministry of Education and Research (BMBF, FKZ 031A216A and B).

## AUTHOR CONTRIBUTIONS

C.C. and I.A. planned and designed the experiments; C.C., performed molecular analyses, phylogenies, expression analysis by RT-qPCR, and transient overexpression of fruits by agro-infiltration; V.M.W developed the 17.66 population and assisted S.L and Y.O in phenotyping. V.W, H.W. and K.O. developed the octoploid SS×FcL mapping population and performed the phenotypic analysis; J.C.T, J. C., Z. L. and I.A. performed the genetic mapping in diploid *F. vesca;* N. O. and P. M. performed fruit quality assessments. R.R., and I.A. performed the genetic mapping and QTL analysis in the octoploid SS×FcL population; C.M-P, D.P. T.T. and T.H. provided genomic data of wild *F. vesca* accessions; N.C., M.A.H. and S.J.K. provided genomic data of cultivated and wild octoploid accessions and participated in KASP marker development; J.G.V. and S.O. conducted the metabolomics analysis; Y.O. C.R.B. and S.L. performed genetic and molecular analyses in UF family 17.66; J.S.S performed RNA-seq and bioinformatic analyses in octoploid strawberry; C.C. and I.A. analyzed the data and wrote the article with inputs from all other authors.

