## Supplemental data for "Allelic Variation of *MYB10* is the Major Force Controlling Natural Variation of Skin and Flesh Color in Strawberry (*Fragaria* spp.) fruit"

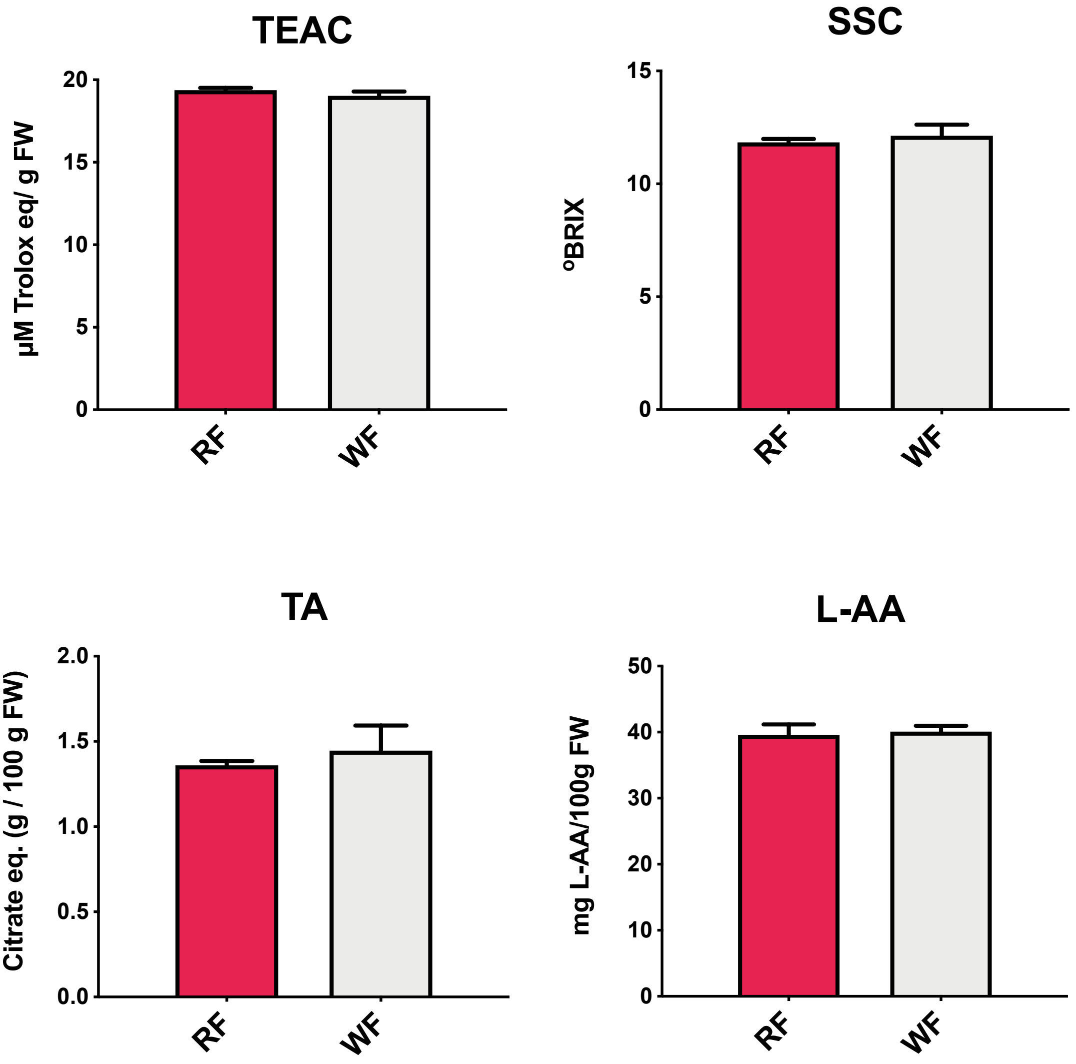


**Supplemental Figure 1.** Total antioxidant capacity (TEAC), soluble solid content (SSC), titratable acidity (TA), and L-ascorbic acid (L-AA) content in RF and WF $F_2$ pools of RV660 x WV596 population.

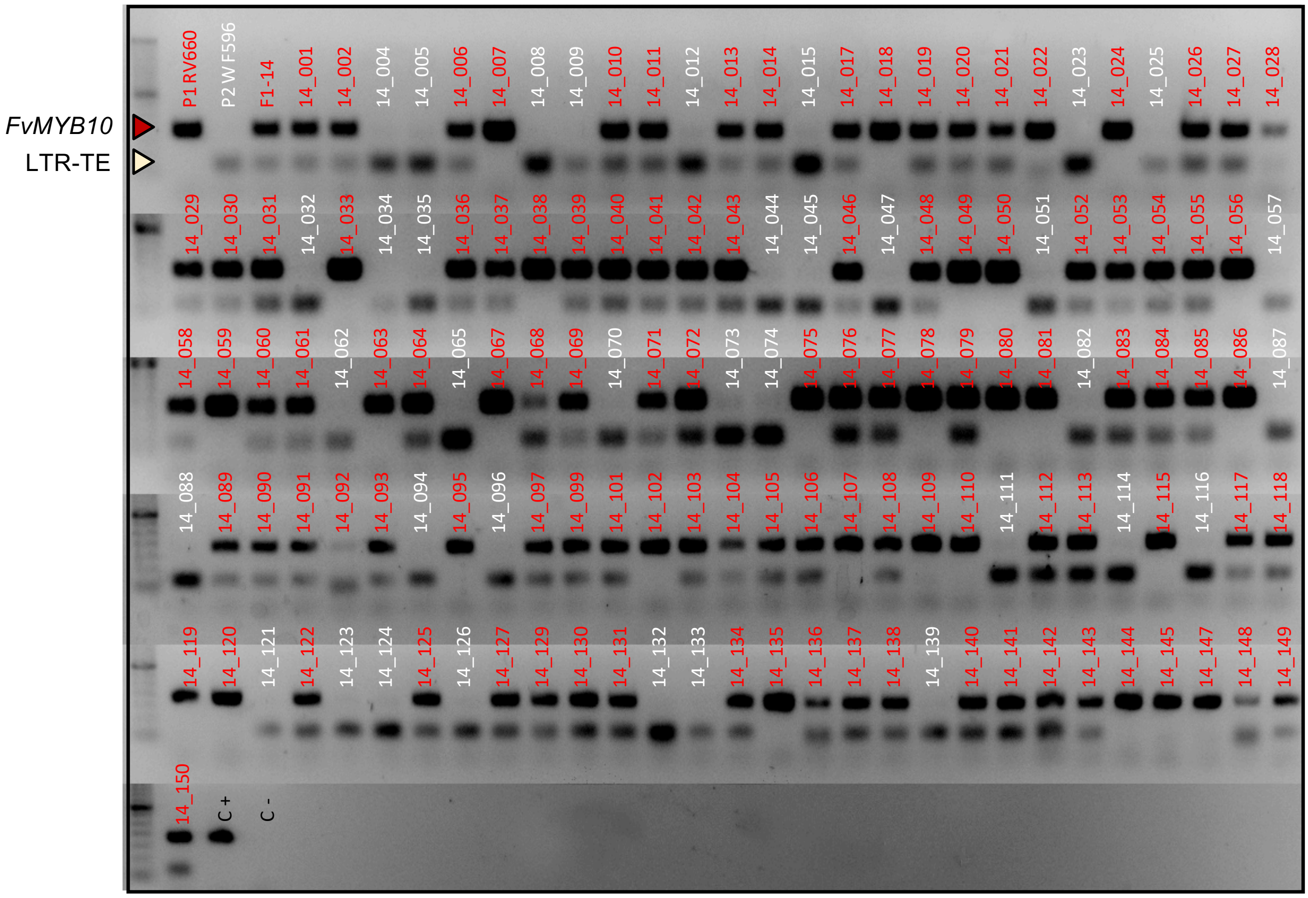


**Supplemental Figure 2.** LTR-TE insertion in *FvMYB10* cosegregates with white fruit phenotype in the entire RV660xWF596 population. PCR was performed using primers R/W-F and R/W-R, flanking the transposon insertion point in *FvMYB10* CDS, together with ccm1, a reverse primer complementary to the 5' end of the 5'LTR (Figure 1C, Supplemental Table 7). Color code in the individual's tag indicates red or white phenotype. All white fruits showed the LTR-TE 59-bp band.

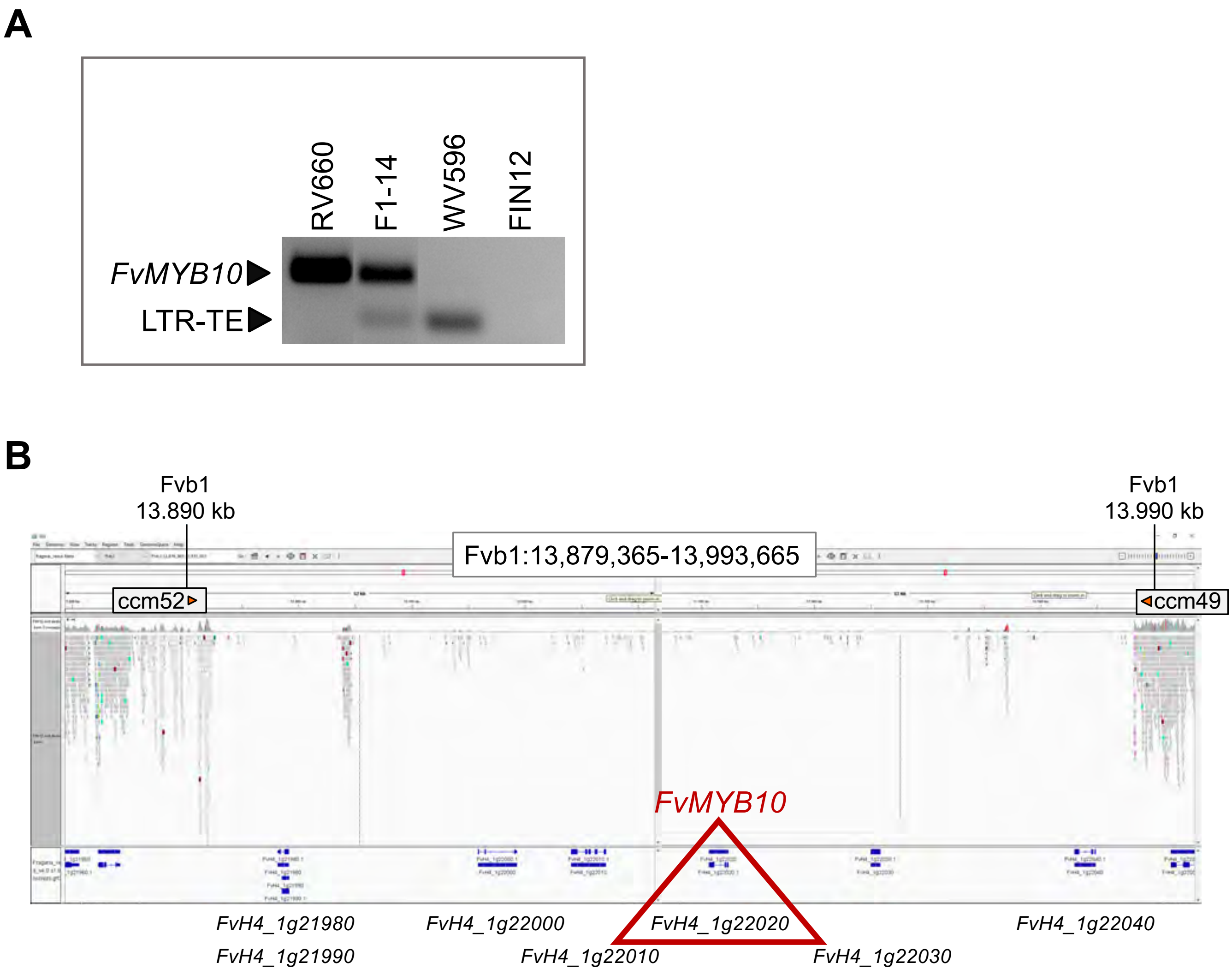


**Supplemental Figure 3.** *F. vesca accession* FIN12 has a large deletion affecting a Fvb1 region of ≈100 kb which results in the loss of 7 predicted genes including *FvMYB10* (*FvH4_1g22020*). (A) PCR with primers R/W-F, R/W-R and ccm1 showing no amplification of FvMYB10 nor the FvMYB10-gypsy LTR-TE from FIN12. (B) Integrative Genomics Viewer (IGV) screenshot showing FIN12 Fvb1 region with almost no sequence coverage. At the bottom, in blue, predicted genes in Hawaii-4 reference genome are shown. Primers ccm52 and ccm49 were used for indel genotyping (Figure 3D).

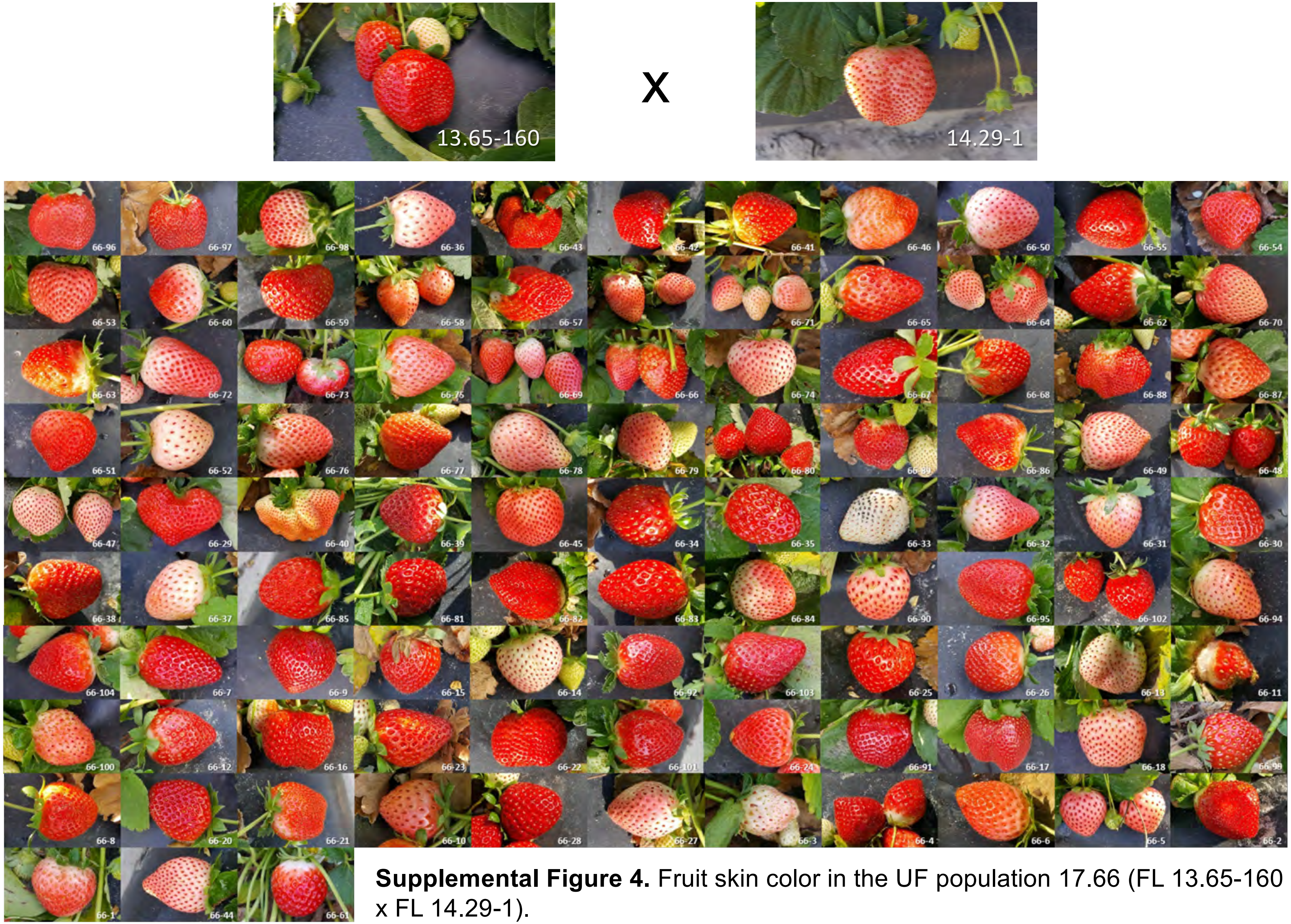


**Supplemental Figure 4.** Fruit skin color in the UF population 17.66 (FL 13.65-160 x FL 14.29-1).

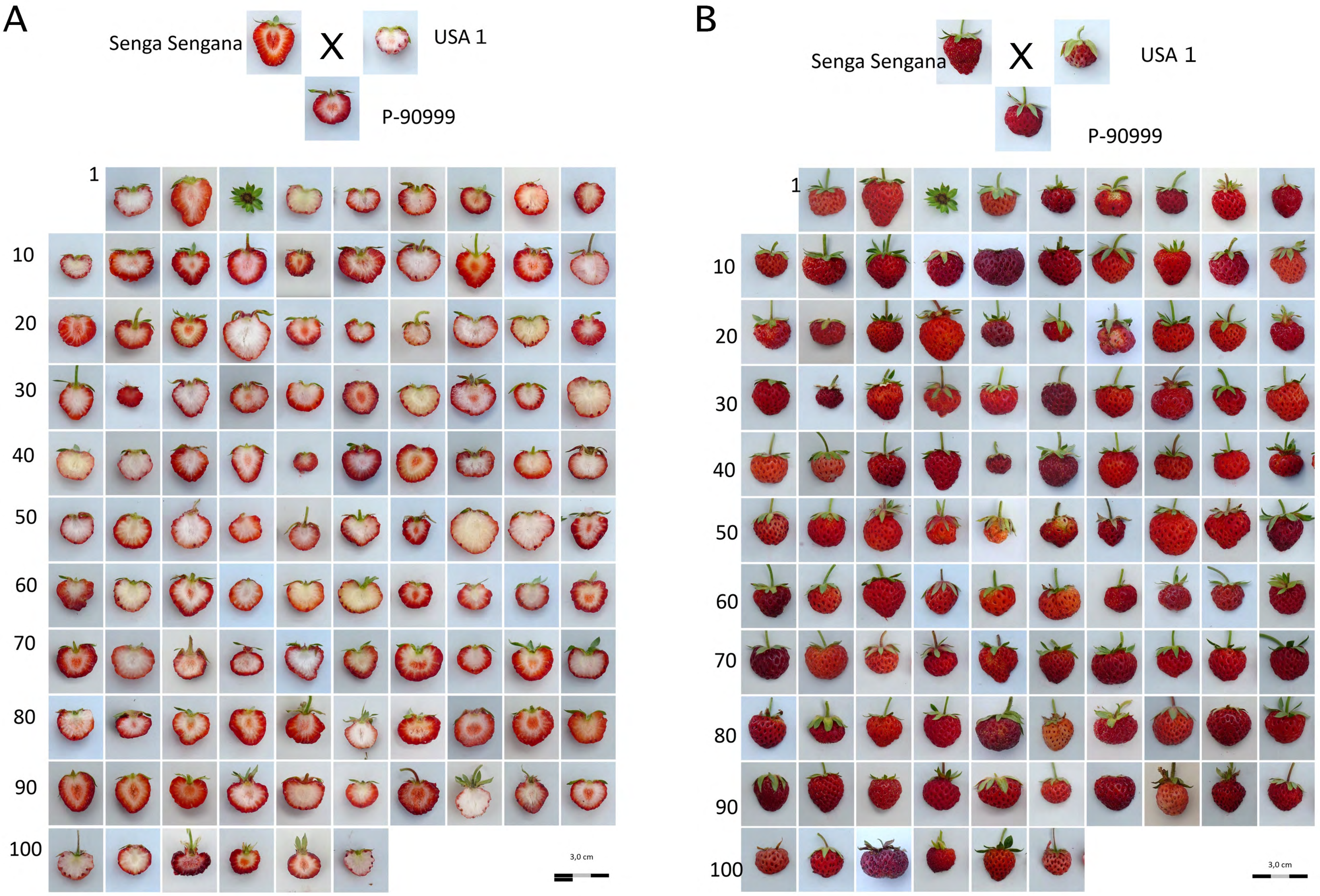


**Supplemental Figure 5.** Fruit flesh (A) and skin (B) color phenotypes in the ‘Senga Sengana’ × *F. chiloensis* ssp. lucida USA2 population. USA 1 as female sister clone of the male USA 2 is shown. P-90999 is the $F_1$ hybrid.

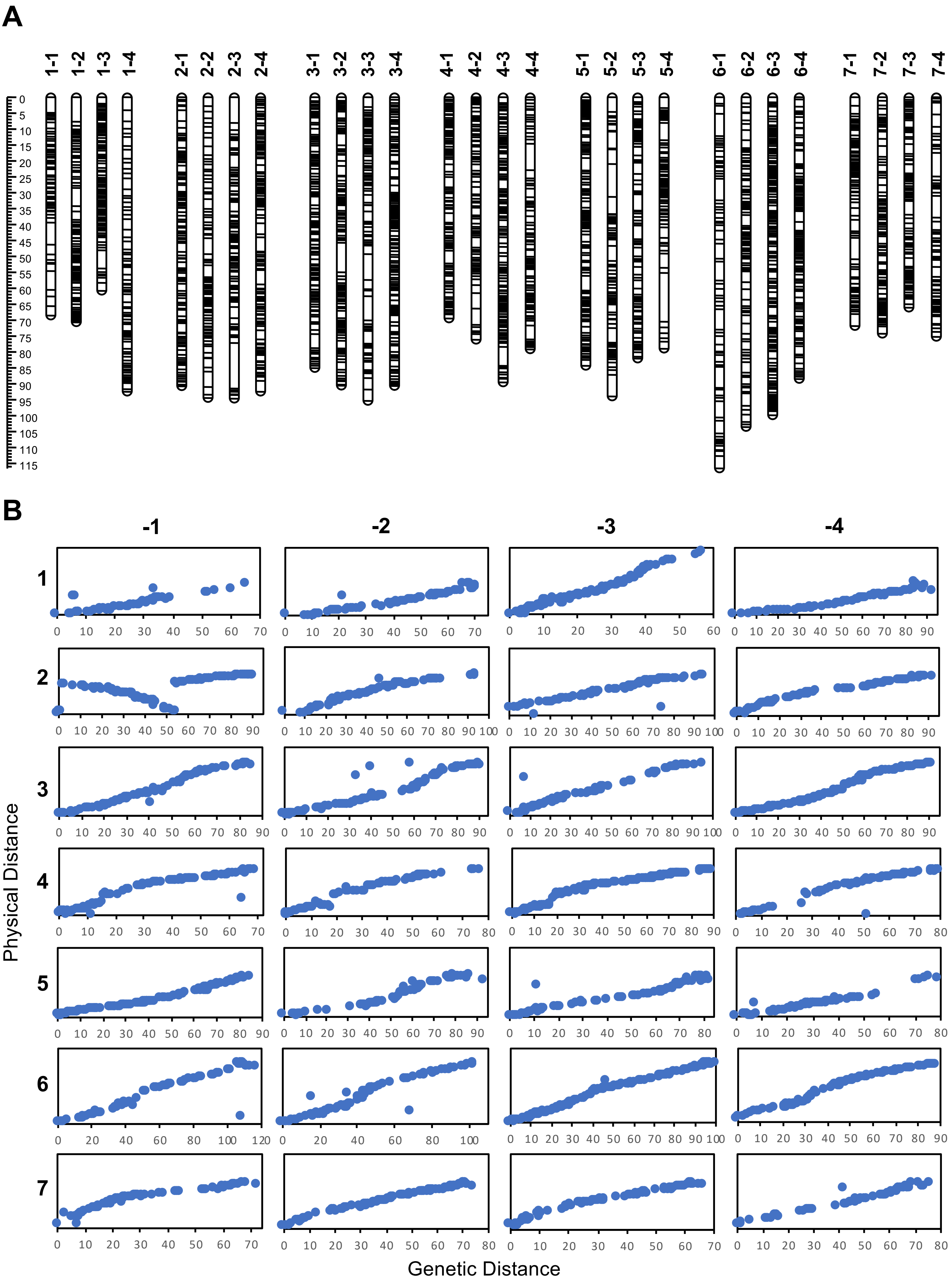


**Supplemental Figure 6. (A)** ‘Senga Sengana’ x *F. chiloensis* ssp. *lucida* linkage map using 2991 DArTseq-derived SNPs. The scale on the left indicates genetic distance (in cM). The black lines in the linkage groups represent the genetic position of markers. LGs within homoeologous groups are named according to the ‘Camarosa’ genome sequence (Edger et al., 2019). **(B)** Marker genetic distances (cM) plotted against *F. vesca* physical positions.

**A**

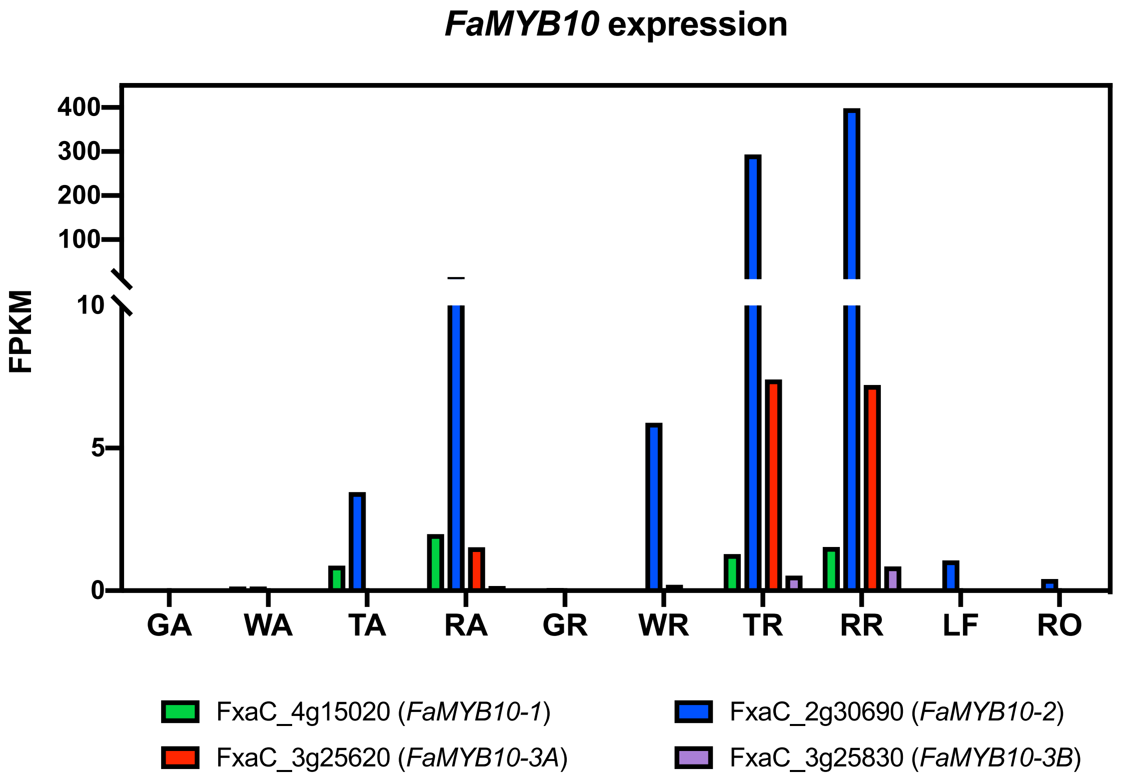


**B**

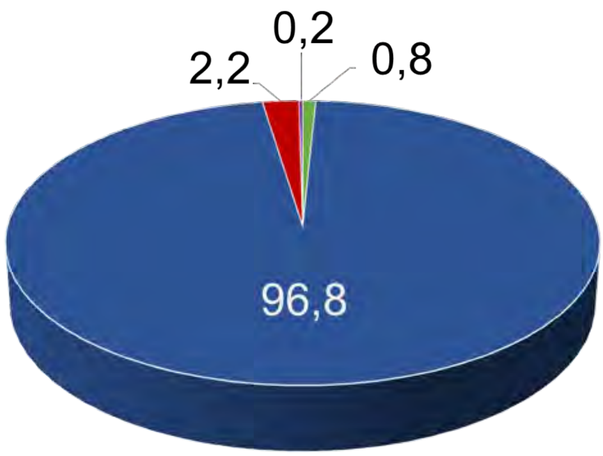


**Supplemental Figure 7.** Expression of *FaMYB10* homoeologs in 'Camarosa'

**(A)** Expression level (FPKM) of all four 'Camarosa' *FaMYB10* homoeologs in achene, receptacle, leaf and root tissues. GA: Green Achene; WA: White Achene; TA: Turning Achene; RA: Red Achene; GR: Green Receptacle; WR: White Receptacle; TR: Turning Receptacle; RR: Red Receptacle; LF: Leaf; RO: Root.

**(B)** Total expression of each *FaMYB10* homoeolog adding up the RFKM values obtained in all tissues analyzed and expressed as a percentage of the global *FaMYB10* expression in the same tissues. Fvb1-2 homoeolog *FxaC_2g30690* (*FaMYB10-2*) is the dominant allele, contributing 96,8% of the expression.

**A**

```
RS    1 ATGGGGGGTTTCGGTGTGAGAAAAGGTGCATGGACTAAAGAGGAAGATGAGCTTCTGAAA
WS    1 ATGGGGGGTTTCGGTGTGAGAAAAGGTGCATGGACTAAAGAGGAAGATGAGCTTCTGAAA

RS   61 CAGTTCATCGAAATCCATGGAGAAGGCAAATGGCATCATGTTCCTCTCAAATCAGGCTTA
WS   61 CAGTTCATCGAAATCCATGGAGAAGGCAAATGGCATCATGTTCCTCTCAAATCAGGCTTA

RS  121 AACAGATGCAGGAAGAGCTGTAGACTGAGGTGGGTGAATTATTTGAAGCCGAATATCAAG
WS  121 AACAGATGCAGGAAGAGCTGTAGACTGAGGTGGGTGAATTATTTGAAGCCGAATATCAAG

RS  181 AGAGGAGAGTTTGCAGAGGATGAAGTTGATTTGATCATCAGGCTTCATAAGCTTCTAGGA
WS  181 AGAGGAGAGTTTGCAGAGGATGAAGTTGATTTGATCATCAGGCTTCATAAGCTTCTAGGA

RS  241 AACAGGTGGTCTTTAATTGCCGGACGATTGCCAGGAAGAACTGCCAATGATGTGAAGAAC
WS  241 AACAGGTGGTCTTTAATTGCCGGACGATTGCCAGGAAGAACTGCCAATGATGTGAAGAAC

RS  301 TATTGGAATACTTATCAAAGGAAAAAGGATCAAAAGACGGCTTCATACGCAAAGAAACTG
WS  301 TATTGGAATACTTATCAAAGGAAAAAGGATCAAAAGACGGCTTCATACGCAAAGAAACTG

RS  361 AAAGTTAAACCCCGAGAAAATACAATAGCTTACACAATTGTAAGACCTCGACCACGAACC
WS  361 AAAGTTAAACCCCGAGAAAATACAATAGCTTACACAATTGTAAGACCTCGACCACGAACC

RS  421 TTCATCAAAAGGTTCAATTTTACAGAGAGAGACGCAAATATAGAGCATAATCATTCAGAA
WS  421 TTCATCAAAAGGTTCAATTTTACAGAGAGAGACGCAAATATAGAGCATAATCATTCAGAA

RS  481 GTGAGTTATAC--------CAGTTCTTTACCAACAGAACCACCACAGACTCTACAATTAG
WS  481 GTGAGTTATACACTTATACCAGTTCTTTACCAACAGAACCACCACAGACTCTACAATTAG

RS  533 AAAATGTAACTGATTGGTGGAAAGATTTCTCAGAAGATAGTACAGAGAGCATTGATAGAA
WS  541 AAAATGTAACTGATTGGTGGAAAGATTTCTCAGAAGATAGTACAGAGAGCATTGATAGAA

RS  593 CAATGTGTTCTGGTCTTGGTTTAGAGGATCATGACTTCTTCACAAACTTTTGGGTTGAAG
WS  601 CAATGTGTTCTGGTCTTGGTTTAGAGGATCATGACTTCTTCACAAACTTTTGGGTTGAAG

RS  653 ATATGGTACTATCGGCAAGCAATCATCTAGTCAACATCTCCTACGTGTGA
WS  661 ATATGGTACTATCGGCAAGCAATCATCTAGTCAACATCTCCTACGTGTGA
```

**B**

```
RS    1 MGGFGVRKGAWTKEEDELLKQFIEIHGEGKWHHVPLKSGLNRCRKSCRLRWVNYLKPNIK
WS    1 MGGFGVRKGAWTKEEDELLKQFIEIHGEGKWHHVPLKSGLNRCRKSCRLRWVNYLKPNIK
        ************************************************************
RS   61 RGEFAEDEVDLIIRLHKLLGNRWSLIAGRLPGRTANDVKNYWNTYQRKKDQKTASYAKKL
WS   61 RGEFAEDEVDLIIRLHKLLGNRWSLIAGRLPGRTANDVKNYWNTYQRKKDQKTASYAKKL
        ************************************************************
RS  121 KVKPRENTIAYTIVRPRPRTFIKRFNFTERDANIEHNHSEVSYTSSLPTEPPQTLQLENV
WS  121 KVKPRENTIAYTIVRPRPRTFIKRFNFTERDANIEHNHSEVSYTLIPVLYQQNHHRLYN*
        ********************************************        :  :* *
RS  181 TDWWKDFSEDSTESIDRTMCSGLGLEDHDFFTNFWVEDMVLSASNHLVNISYV---
WS  180 KM*LIGGKISQKIVQRALIEQCVLVLV*RIM[illegible]SSQTFGLKIWYYRQAII*[illegible]TSPTC
        .    . . ...      : . : :    ::*.  . .: :   .: :  *
```

**Supplemental Figure 8**. Sequence alignment of WT and mutant MYB10-2 in the UF breeding population 17.66

(**A**) Genomic DNA sequence alignment of *MYB10-2* coding region of red and white strawberry. RS and WS represent red and white strawberry, respectively. Conserved sequences are highlighted in black. Deletion region labelled as '-'.

(**B**) An amino-acid sequence alignment of predicted MYB10-2 of red and white strawberry. RS and WS represent red and white strawberry, respectively. Residues that are conserved across all sequences are highlighted in black. Below the protein sequences is a key denoting conserved sequence (*), conservatie mutations (:), semi-conservative mutations (.), and non-conservative mutations ( ).

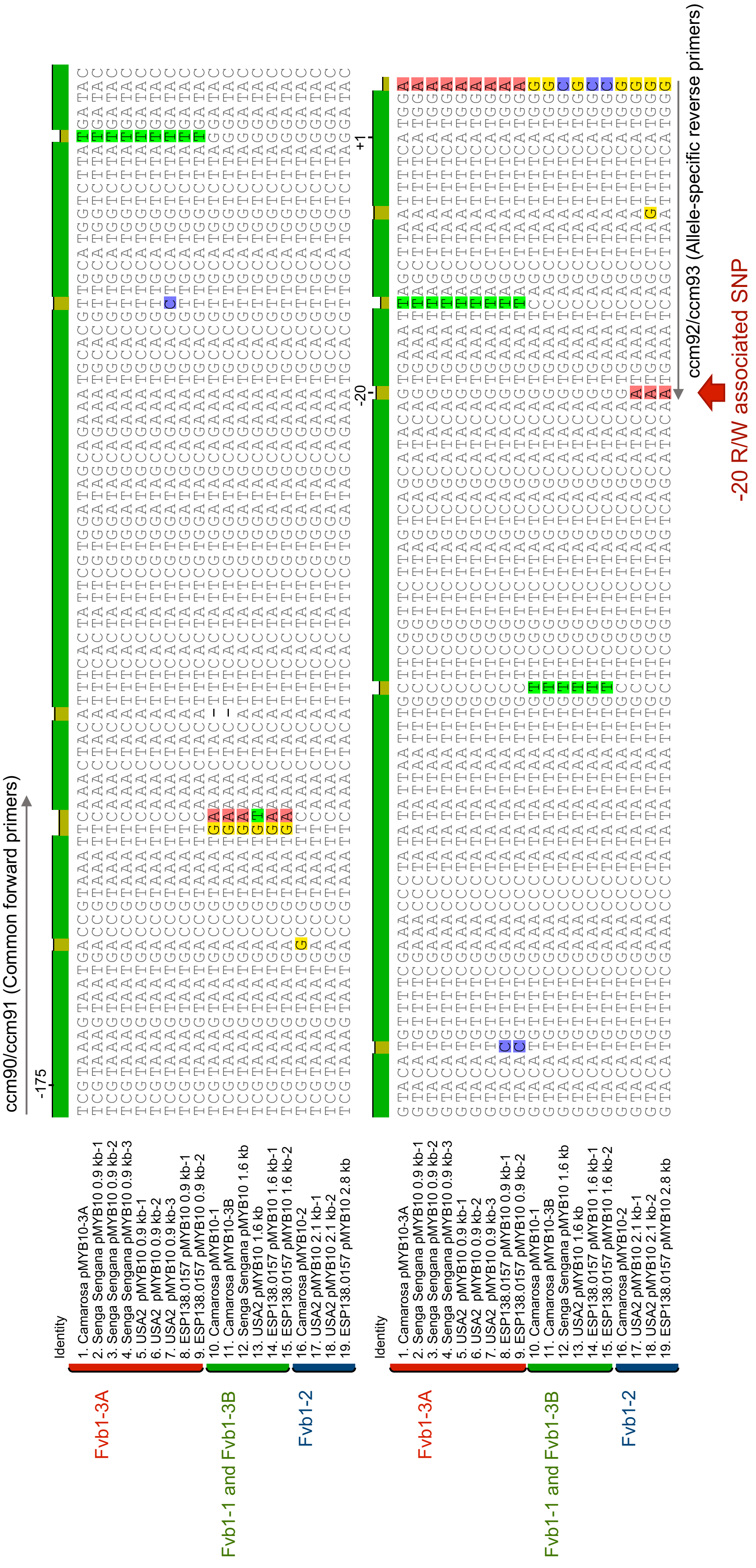


**Supplemental Figure 9.** *MYB10* upstream region alignment showing the SNP targeted for KASP genotyping.

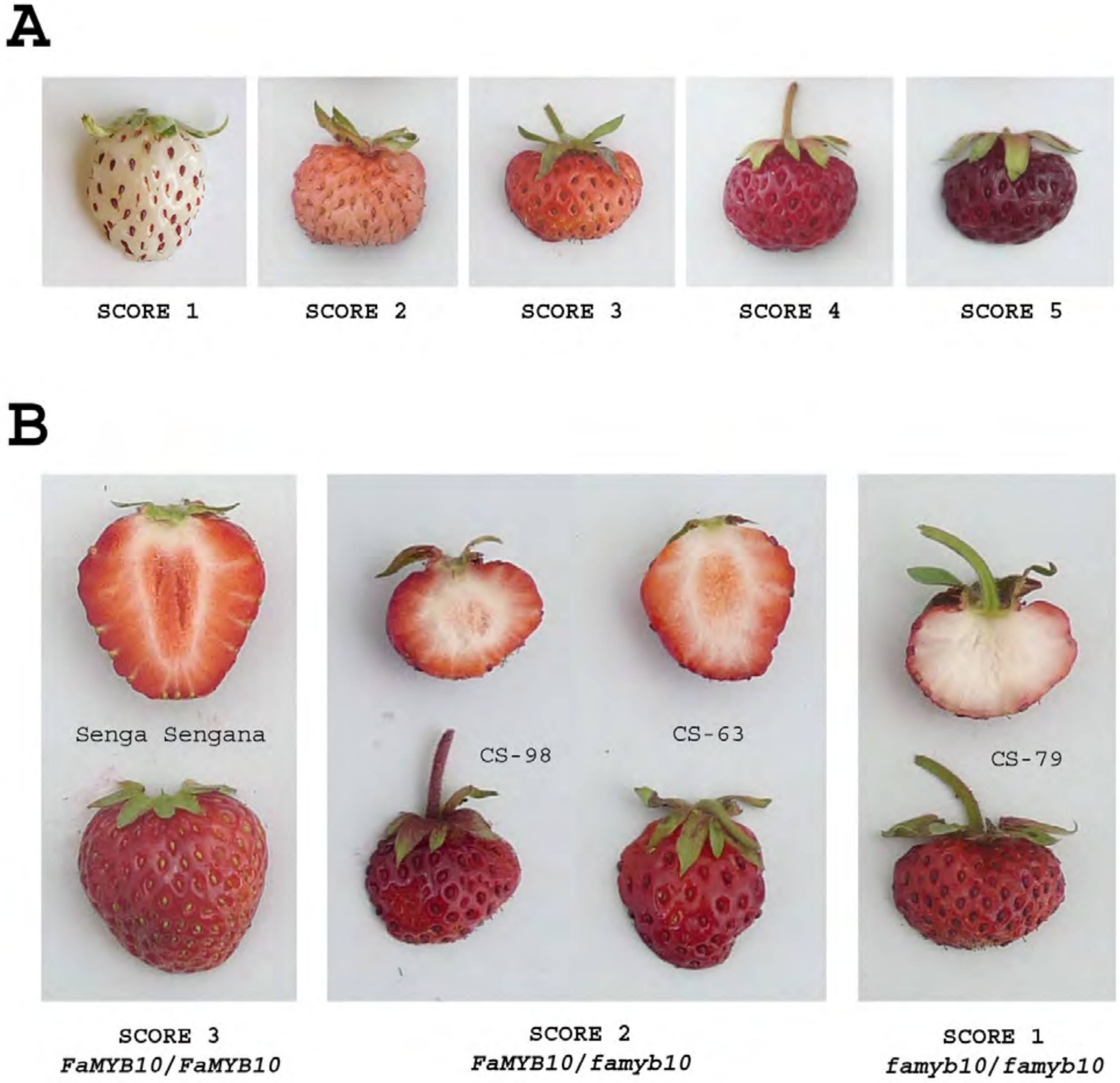


**Supplemental Figure 10.** Fruit color scale used to phenotype octoploid accessions. (**A**) Skin color scale (1-5). (**B**) Flesh color scale (1-3). Examples used in B had a score 4 in skin color. CS-63, CS-79 and CS-98 are different $F_2$ lines from Hansabred SS×FcL cross.
